# PSF-Driven Spatio-Temporal Blending in Fluorescence Lifetime Imaging Microscopy and Its Mitigation via Mean-Shift Super-Resolution-Based Masking

**DOI:** 10.64898/2026.03.17.712453

**Authors:** Mario González-Gutiérrez, Diana M. Vázquez-Enciso, Nicolás Mateos, Wonsang Hwang, Esley Torres-García, Haydee O. Hernández, Jenu V. Chacko, Iván Coto Hernández, Pablo Loza-Álvarez, Christopher Wood, Adan Guerrero

## Abstract

Fluorescence Lifetime Imaging Microscopy (FLIM) enables quantitative mapping of molecular environments in living systems with high biochemical specificity. However, spatial overlap dictated by the diffraction-limited point spread function (PSF) causes a mixing of temporal signals: photons from neighboring emitters collected within the same pixel yield composite decay profiles, generating apparent intermediate lifetimes that can be mistaken for variations in the local molecular environment. We introduce a workflow that applies Mean-Shift Super-Resolution (MSSR) to raw intensity data to generate intensity-derived spatial masks prior to phasor-based lifetime analysis. The method is computationally efficient and preserves decay kinetics because it operates on intensity-derived spatial information rather than modifying temporal data. In U2OS cells labeled with spectrally-overlapping fluorophores, phasor analysis reveals an intermediate lifetime population localized at PSF-overlap interfaces, consistent with optical mixing rather than intrinsic lifetime heterogeneity. MSSR-derived masking suppressed this mixed population while preserving stable phasor cluster centers –*i.e*. the distribution of similar phasor coordinates in the phasor plane– for each fluorophore. Simulations of strictly monoexponential fluorescence decay emitters further show that blended lifetime decay profiles are present at separations up to 4σ and becomes maximal near ∼1.6σ, indicating that conventional spatial resolution criteria can underestimate lifetime cross-talk. Application of this workflow to three-component FLIM showed also a reduced overlap of pixel distributions in phasor plots while maintaining distinct lifetime signatures. Overall, MSSR-based spatial refinement provides an accessible strategy to improve the spatial resolution while maintaining accuracy of FLIM measurements.

## INTRODUCTION

Fluorescence Lifetime Imaging Microscopy (FLIM) maps the spatial distribution of fluorescence decay kinetics and thereby reports on biochemical and biophysical states that are often inaccessible to intensity-only imaging. By probing excited-state relaxation dynamics, FLIM enables contrast based on environmental sensitivity and molecular interactions even when fluorophore concentration, excitation inhomogeneity, or optical path variations confound intensity measurements. FLIM implementations are commonly classified into time-domain (TD-FLIM) and frequency-domain (FD-FLIM) modalities. [1–5] TD-FLIM estimates lifetimes from the distribution of photon arrival times after pulsed excitation, most often using time-correlated single-photon counting (TCSPC) with single-photon detectors (including photomultiplier-based and single-photon avalanche diode (SPAD)-based detectors) or using widefield time-gated detection, and recent advances include analog approaches that record fluorescence waveforms. [1–5] FD-FLIM, in contrast, derives lifetimes from the phase shift and demodulation of fluorescence emission signal relative to a temporally modulated excitation signal. [5] FLIM has found extensive applications in medical imaging and in biochemical and physicochemical research, enabling quantitative assessments of parameters such as pH, viscosity, and other environmental or material properties. [3,6-17]

In FLIM measurements, uncertainty in the temporal characterization of fluorescence decays can arise from several distinct sources. First, the instrument response function (IRF) limits the temporal resolution with which photon arrival times can be resolved, imposing an instrumental broadening of the measured decay. Second, the stochastic nature of photon detection introduces shot-noise–limited uncertainty, particularly under low-count regimes. Third, and central to the present work, diffraction-limited imaging induces spatial convolution through the optical point spread function (PSF), such that photons emitted by neighboring fluorophores are collected within the same pixel. This spatial overlap produces composite decay signals that manifest as apparent lifetime mixtures that do not correspond to intrinsic photophysical interactions.[1–5] We refer to this effect as PSF-induced Temporal Blending (TB).

As a historical note, the impact of spatial overlap on lifetime measurements has been recognized in early FLIM studies. To the best of our knowledge, the first explicit description of this phenomenon was provided by Philippe Bastiaens and Anthony Squire, who introduced the term Temporal Blurring.[18] While this terminology effectively conveys the decrease of precision of the FLIM measurement, in the present work we adopt the term Temporal Blending to reflect more precisely the underlying mechanism: the mixing of photon streams from spatially overlapping PSFs prior to lifetime transformation. In this framework, the measured decay represents a spatially weighted superposition of multiple fluorescence decays within the diffraction-limited observation volume. Importantly, TB can mimic genuine lifetime heterogeneity arising from environmental variations, molecular interactions, or conformational dynamics, thereby complicating the interpretation of FLIM measurements. [18,19]

Furthermore, it is important to distinguish this effect from fundamentally different phenomena such as diffraction in time[20]. While PSF-induced temporal blending represents a classical, instrumentally induced effect arising from optical diffraction, diffraction in time is a quantum-mechanical phenomenon associated with the time–energy uncertainty principle and describes the transient behavior of massive particles governed by the Schrödinger equation when encountering temporal boundaries[20]. Despite the similarity in terminology, this effect is unrelated to the optical measurement process considered here.

Independent of acquisition modality, lifetime estimation can be performed using fitting or non-fitting approaches. Fitting methods include weighted least squares estimation (WLS), maximum likelihood estimation (MLE), Global and Bayesian fitting. Non-fitting routines include rapid lifetime determination (RLD), Center-of-mass methods (CMM), Integral extraction methods, Laguerre Expansion and the Phasor approach. [21] More recently, machine learning and deep learning strategies have been explored to improve estimation accuracy and throughput, and to provide denoising capabilities. [22-25]

The phasor method provides a graphical representation of fluorescence decay kinetics by mapping each pixel to phasor coordinates (G,S). In this framework, monoexponential decays lie on the universal semicircle, which represents the geometric locus of phasor coordinates corresponding to lifetimes ranging from zero to infinity (for a given modulation frequency). Multiexponential decays fall inside this semicircle, where their position corresponds to the vector sum of the phasors of the individual lifetime components, weighted by their relative contributions. Intermediate phasor positions can reflect either genuine biochemical heterogeneity within the sampled volume or linear combinations of distinct lifetime components. This mixing property is useful for unmixing and segmentation, but it also implies that intermediate lifetimes can arise from optical cross-talk when photons from multiple emitters are pooled within a single spatial bin. [26-30]

As introduced above, PSF-induced Temporal Blending leads to spatially mixed decay signals that can bias lifetime estimation in diffraction-limited conditions. A direct strategy to mitigate TB is to increase spatial resolution so that neighboring emitters are better separated prior to lifetime extraction. Super-resolution techniques can be broadly classified into three categories. The first comprises deterministic approaches such as Stimulated Emission Depletion (STED) and Structured Illumination Microscopy (SIM). [31–36] STED has been integrated with lifetime measurements through multiple strategies, including time-gated STED, Separation of Photons by Lifetime Tuning (SPLIT-STED), STED-FLIM combined with phasor analysis, and TauSTED, which exploit depletion-induced lifetime gradients to refine contrast and spatial information. [37–44] The second category comprises stochastic localization-based super-resolution methods collectively referred to as Single-Molecule Localization Microscopy (SMLM). Representative SMLM implementations include Photoactivated Localization Microscopy (PALM) and Stochastic Optical Reconstruction Microscopy (STORM), which reconstruct super-resolved images by localizing temporally separated single-emitter signals over many acquisition frames. [45-47] Lifetime-resolved variants, including confocal fluorescence-lifetime SMLM and time-correlated SMLM, further increase molecular specificity by assigning lifetime values to localized molecules using TCSPC-based detection. [48,49]

The third category comprises computational approaches, including deconvolution methods that attempt to reverse optical blurring using an estimated PSF, as well as fluctuation-based super-resolution methods such as XC-SOFI, 3B, ESI, and SRRF. [47] Addressing TB specifically, Bastiaens and Squire developed an iterative constrained deconvolution method based on the Tikhonov–Miller algorithm (ICTM), processing temporal information through Fourier components that are trigonometric analogs of the phasor coordinates (G,S). [18,50] While effective in relevant contexts, many computational super-resolution strategies require large frame stacks, substantial computation, or temporal processing steps that may introduce artifacts or confounds when applied to FLIM, particularly when preservation of decay kinetics is essential. [47]

Mean-Shift Super-Resolution (MSSR) offers a distinct opportunity for FLIM because it operates on intensity information to enhance spatial separability without directly altering the fluorescence decay kinetics. [51] In the present workflow, MSSR is applied to an average (or integrated) intensity projection of the FLIM dataset to generate an intensity-derived spatial mask with improved separability and signal-to-noise ratio. Phasor coordinates are then computed from the unmodified time-resolved FLIM data within the masked regions, thereby suppressing PSF-driven photon mixing while conserving lifetime signatures. Accordingly, we develop and validate a method that couples MSSR-derived spatial masks with phasor analysis, demonstrating that improved spatial separability can reduce Temporal Blending in FLIM while maintaining decay fidelity in complex specimens.

## RESULTS

### Spatial convolution drives temporal blurring in lifetime imaging

To experimentally illustrate how Temporal Blending (TB) emerges at the cellular scale, we analyzed a fixed-cell specimen in which two distinct subcellular structures—mitochondria and microtubules—were labeled with spectrally overlapping fluorophores. Alexa Fluor 532 (AF532; TOM20) and Alexa Fluor 555 (AF555; α-tubulin) have closely spaced emission maxima (553 nm and 580 nm), rendering the two structures indistinguishable in single-channel confocal intensity imaging (Fig. 1a). In this reference specimen, both dyes are immobilized via immunolabeling on their respective targets, and no direct biochemical coupling between labels is expected. The system therefore approximates two independent lifetime species; under this condition, intermediate apparent lifetimes are predicted to arise predominantly from optical photon pooling when diffraction-limited point spread functions (PSFs) overlap.

**Fig. 1.**
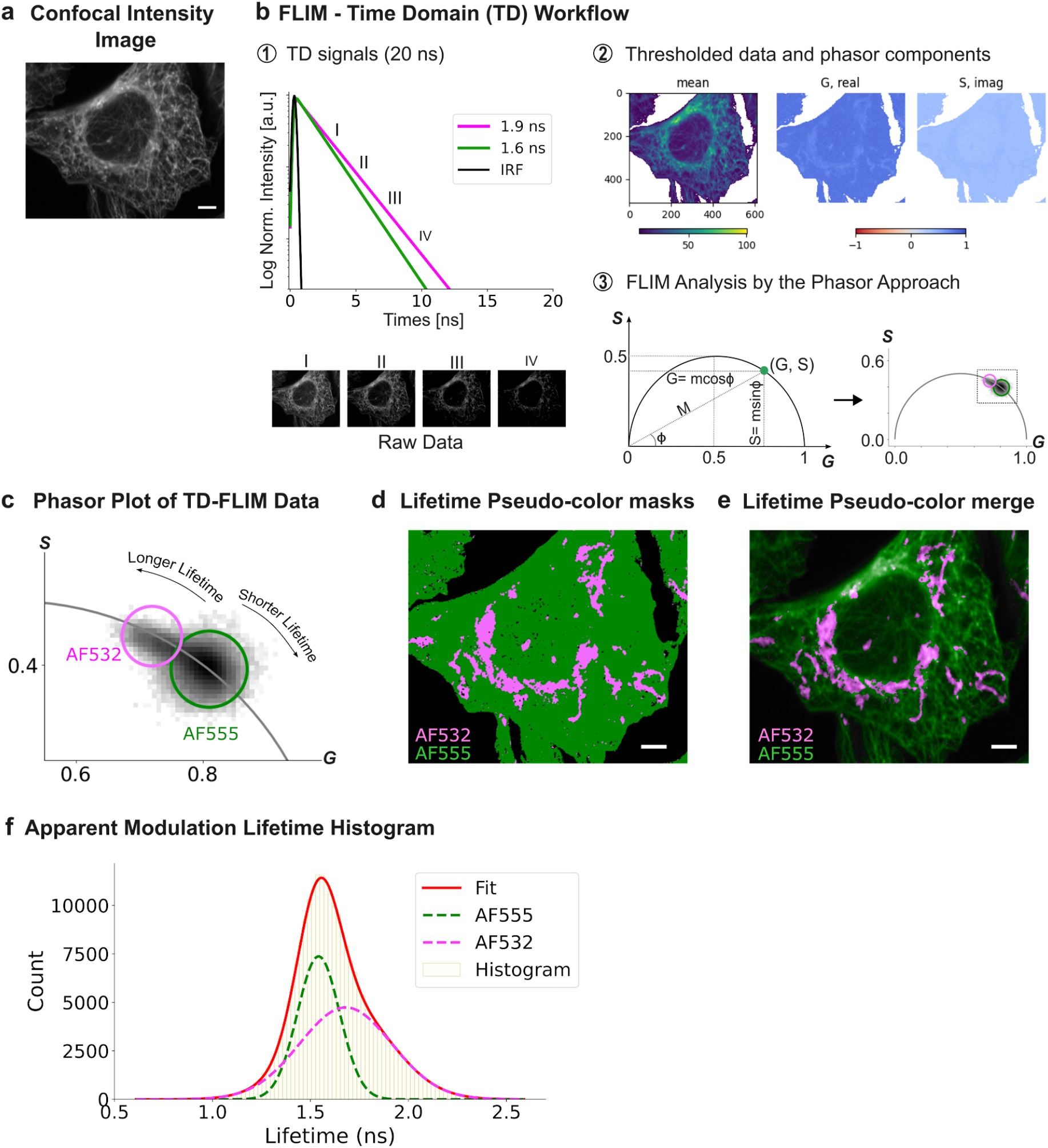
TD-FLIM phasor workflow and endpoint lifetime populations in a dual-labelled cellular specimen. (a) Single-channel confocal intensity image of a U2OS cell specimen labeled for mitochondria (AF532; TOM20) and microtubules (AF555; α-tubulin); structures are not separable in the intensity domain due to spectral proximity. (b) TD-FLIM workflow schematic. (b1) Each pixel contains a time-resolved intensity profile; schematic monoexponential decays illustrate two independent pixels dominated by different fluorophore species (short- and long-lifetime exemplars), corresponding to the qualitative endpoint behaviors expected for AF555- and AF532-enriched regions in this specimen. Traces are schematic and included for conceptual reference rather than dye-specific calibration. (b2) Mean intensity image and phasor-component images (G and S) after intensity thresholding. (b3) Phasor representation schematic: a single-component decay maps to a point on the universal semicircle (left), whereas the experimental data exhibit two lobes (right), displayed in detail in (c). (c) Phasor plot of experimental TD-FLIM data with two endpoint cursor selections defining AF532-enriched and AF555-enriched populations. (d) Pseudo-color lifetime mask obtained by projecting cursor selections from (c) into image space. (e) Pseudo-color mask merged with intensity. (f) Apparent modulation lifetime histogram for endpoint-selected populations with Gaussian fits. Scale bar: 2.7 μm.

Because the core claim evaluated here is that intermediate apparent lifetimes can arise from optical mixing, we first establish the relevant phasor geometry and the relationship between time-domain (TD) and frequency-domain (FD) FLIM using a didactic overview (Supporting Information, Fig. S1). TD-FLIM samples fluorescence relaxation as a time-resolved intensity decay following pulsed excitation (Fig. S1a), whereas FD-FLIM derives lifetimes from the phase delay and demodulation of the emission waveform relative to sinusoidally modulated excitation (Fig. S1b). Both modalities yield equivalent phasor coordinates (G,S) for the same decay at a given harmonic, taking into account calibration and instrument-response corrections; consequently, TD- and FD-FLIM provide indistinguishable phasor plots in the ideal case[30] (Fig. S1c). In this representation, monoexponential decays map to single points on the universal semicircle whereas multicomponent decays and linearly-mixed signals map inside the semicircle [26-30](Fig. S1c). Importantly, a pixel receiving photons from distinct species of monoexponential emitters yields an amplitude-weighted composite decay whose phasor lies between the endpoint phasors according to the same linear combination rule. This framework defines the operational signature of TB: intermediate phasor positions that arise from spatial photon pooling rather than from lifetime heterogeneity due to environmental parameters or molecular interactions.

With this foundation, Fig. 1 summarizes the TD-FLIM phasor workflow used to evaluate TB in the cellular specimen. Each pixel contains a temporal-domain intensity profile sampled across time gates, and the full set of pixel-wise decays is analyzed to compute phasor-component images (G and S) and a mean intensity image after intensity thresholding (Fig. 1b2). Fig. 1b1 shows schematic monoexponential decay exemplars for two independent pixels dominated by different fluorophore species, illustrating the endpoint behaviors expected for a two-species specimen (schematic traces are included for conceptual reference rather than dye-specific calibration under the present conditions). Phasor coordinates derived from all pixels populate the phasor plot (Fig. 1b3,c), and the reciprocity between phasor space and image space enables bidirectional mapping: cursor selections in phasor space can be projected back into the image to generate spatial masks, and spatial regions of interest can be mapped to their corresponding phasor distributions.

The experimental phasor distribution exhibited two dominant lifetime populations (Fig. 1c). Rather than forming two fully separated compact clusters, the data displayed two partially separated lobes connected by an elongated continuum between the endpoint regions, consistent with pixels containing composite decays. Two circular cursors were placed at the ends of the distribution to define endpoint populations enriched in the short- and long-lifetime components. Projecting these endpoint selections back into image space generated pseudo-color lifetime masks (Fig. 1d) that segmented microtubules predominantly into the short-lifetime class (green; AF555-enriched) and mitochondria predominantly into the long-lifetime class (pink; AF532-enriched). The merged pseudo-color/intensity representation (Fig. 1e) preserved this spatial organization across the cell while highlighting interface zones where class separation became locally heterogeneous, consistent with spatial mixing.

To quantify the endpoint populations, apparent modulation lifetime histograms were computed from the pixels selected by the endpoint cursors (Fig. 1f). The histogram was described by two main components corresponding to the endpoint classes. The short-lifetime endpoint population (AF555-enriched; green) appeared comparatively narrow, whereas the long-lifetime endpoint population (AF532-enriched; pink) was broader, indicating increased dispersion in apparent lifetimes within the long-lifetime class. This broadening suggested that a fraction of pixels assigned to the endpoint classes still contained mixed contributions, motivating explicit identification of intermediate phasor positions expected under the optical mixing behavior introduced in Fig. S1.

To localize intermediate apparent lifetimes, a third cursor was placed on the central region between the endpoint lobes (Fig. 2a). This selection isolated pixels whose phasor coordinates fell between the endpoint populations, consistent with composite decays. Mapping these intermediate pixels back into image space revealed preferential localization at interfaces between mitochondria and microtubules (Fig. 2b), where PSF overlap and photon pooling from adjacent emitters are expected to be maximal. The merged pseudo-color representation (Fig. 2c) further emphasized that the intermediate class (yellow; TB) was spatially confined and enriched at boundary and crossing regions rather than uniformly distributed within either structure. Representative regions of interest (ROI 1–3) highlight the characteristic spatial motif of TB pixels decorating mitochondria–microtubule appositions.

**Fig. 2.**
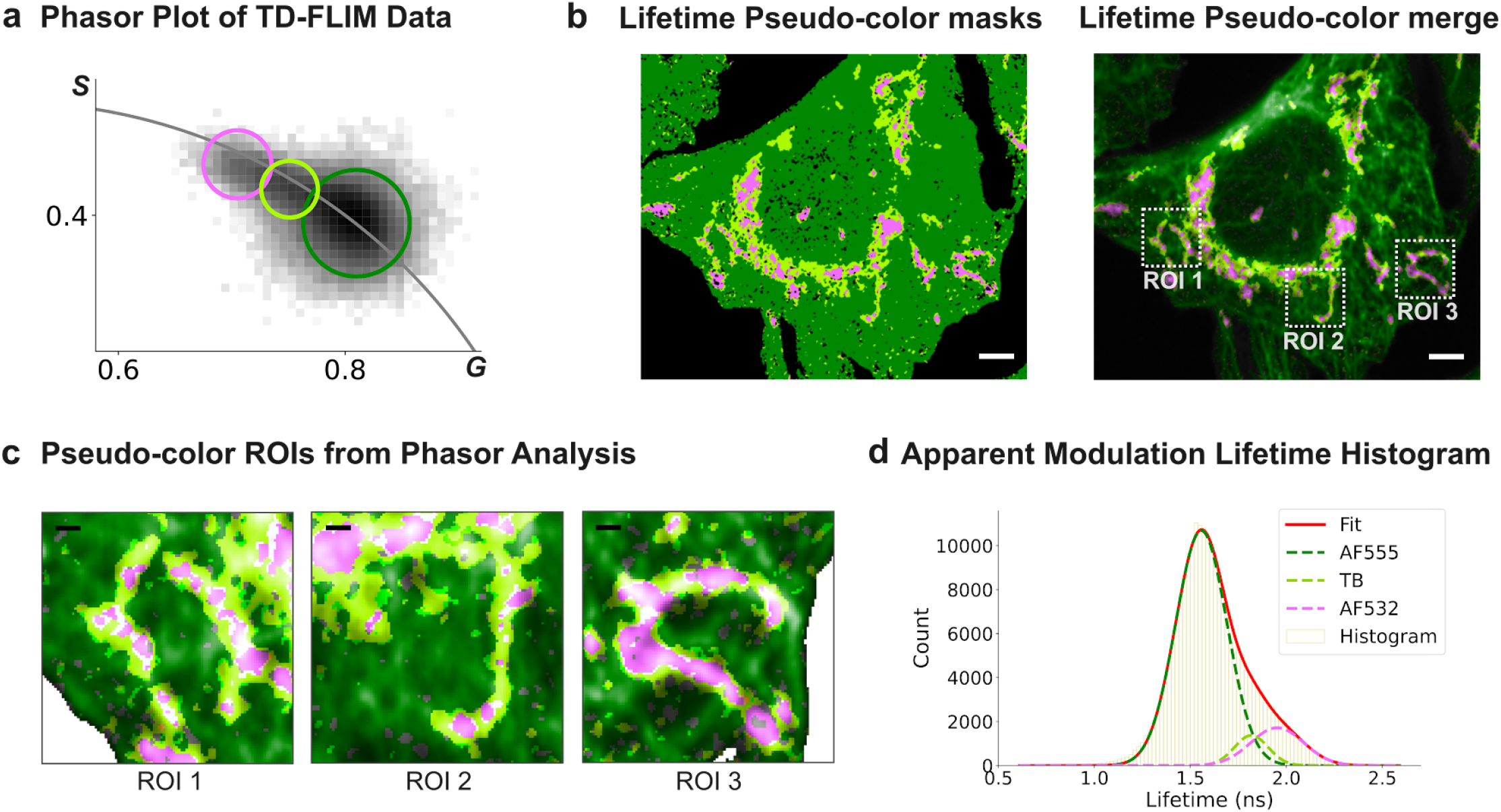
Temporal blending manifests as an intermediate phasor population localized at mitochondria–microtubule interfaces. (a) Phasor plot showing endpoint populations (AF532-enriched, pink; AF555-enriched, green) and an intermediate population (yellow) selected by a third cursor positioned between endpoint lobes. (b) Pseudo-color segmentation map obtained by projecting the three cursor selections into image space, highlighting the intermediate population at structural interfaces. (c) Pseudo-color merge with intensity overlay and three representative regions of interest (ROI 1–3) illustrating interface-localized intermediate pixels. (d) Apparent modulation lifetime histogram computed from the three pixel classes with a triple-Gaussian fit separating AF555-enriched, AF532-enriched, and TB-associated intermediate components. Scale bars in (b): 3 μm; scale bars in (c): 500 nm.

Quantitatively, adding the intermediate pixel class revealed an apparent modulation lifetime distribution that was best described by three components (Fig. 2d): a short-lifetime endpoint (AF555-enriched), a long-lifetime endpoint (AF532-enriched), and an intermediate component assigned to TB. Because both fluorophores are immobilized by immunolabeling on distinct, non-coupled targets (α-tubulin versus TOM20), a third interaction-derived lifetime population (for example, FRET or chemically-coupled photophysics) is not expected at the scale of these structures. Instead, an intermediate apparent lifetime is predicted where diffraction-limited sampling volumes pool photons from both labels, yielding an amplitude-weighted composite decay that falls between endpoints in phasor space. The phasor continuum linking endpoint lobes (Fig. 1c), the interface-enriched localization of intermediate pixels (Fig. 2b,c), and the intermediate peak resolved by three-component fitting (Fig. 2d) together support PSF-driven photon mixing as the source of spatially patterned intermediate lifetimes, establishing a baseline for evaluating masking strategies that improve spatial separability while preserving endpoint signatures.

### Mean-Shift Super-Resolution mitigates Temporal Blending in Fluorescence Lifetime Imaging Microscopy

The baseline Temporal Blending (TB) signature established above, defined as a spatially patterned intermediate lifetime class concentrated at structural interfaces, lead us to explore a strategy that increases spatial separability without direct manipulation of decay kinetics. Mean-Shift Super-Resolution (MSSR) operates on the spatial intensity landscape and can reduce effective PSF overlap through redistribution of signal density toward local maxima, thereby improving emitter separability in diffraction-limited data [47,51-52]. In the present workflow, MSSR is used as a spatial prior that defines where photons are likely to originate, after which lifetime information is interpreted from the original time-resolved measurements. This ordering is essential: spatial processing is applied to an intensity projection to derive a mask, and phasor coordinates remain derived from the unmodified time-domain FLIM data, preserving the decay information that underpins the phasor representation.

A practical aspect of this implementation is that the masking step uses an intermediate bounded output from MSSR rather than the final remapped intensity image. Specifically, the procedure intentionally omits the last MSSR intensity remapping (normalization step) and instead uses the intermediate matrix that is naturally bounded between 0 and 1, denoted here as 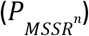. This quantity behaves as a probability-like spatial support map, *i*.*e*. values near 1 concentrate around likely emitter locations, whereas low values occupy background and overlap regions. Thresholding 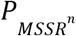 therefore yields binary masks that progressively restrict analysis to spatial regions with high localization likelihood, a property exploited here to mitigate mixed-pixel signatures while maintaining the endpoint lifetime classes identified previously.

The expected effect of MSSR on separability is illustrated using a two-emitter model at the Sparrow limit (two Gaussian emitters separated by 2*σ*). In the raw summed profile, strong overlap yields a single broadened maximum and poor resolvability at the midpoint (Fig. 3a). After MSSR processing, the probability-like output becomes increasingly bimodal as MSSR order increases, enabling threshold selection that isolates high-support cores of each emitter while suppressing the overlap region (Fig. 3b). The corresponding two-dimensional maps illustrate that, across threshold levels, Raw data retain substantial overlap, whereas MSSR^0^ and MSSR^1^ progressively concentrate probability density into separated lobes (Fig. 3c). These model results define the operational principle used in the cellular experiment. Fig. 4 applies the same 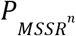-thresholding logic motivated in Fig. 3 to generate a spatial mask from a FLIM intensity projection, with the goal of suppressing overlap-dominated pixels that are predicted to underlie intermediate phasor positions.

**Fig. 3.**
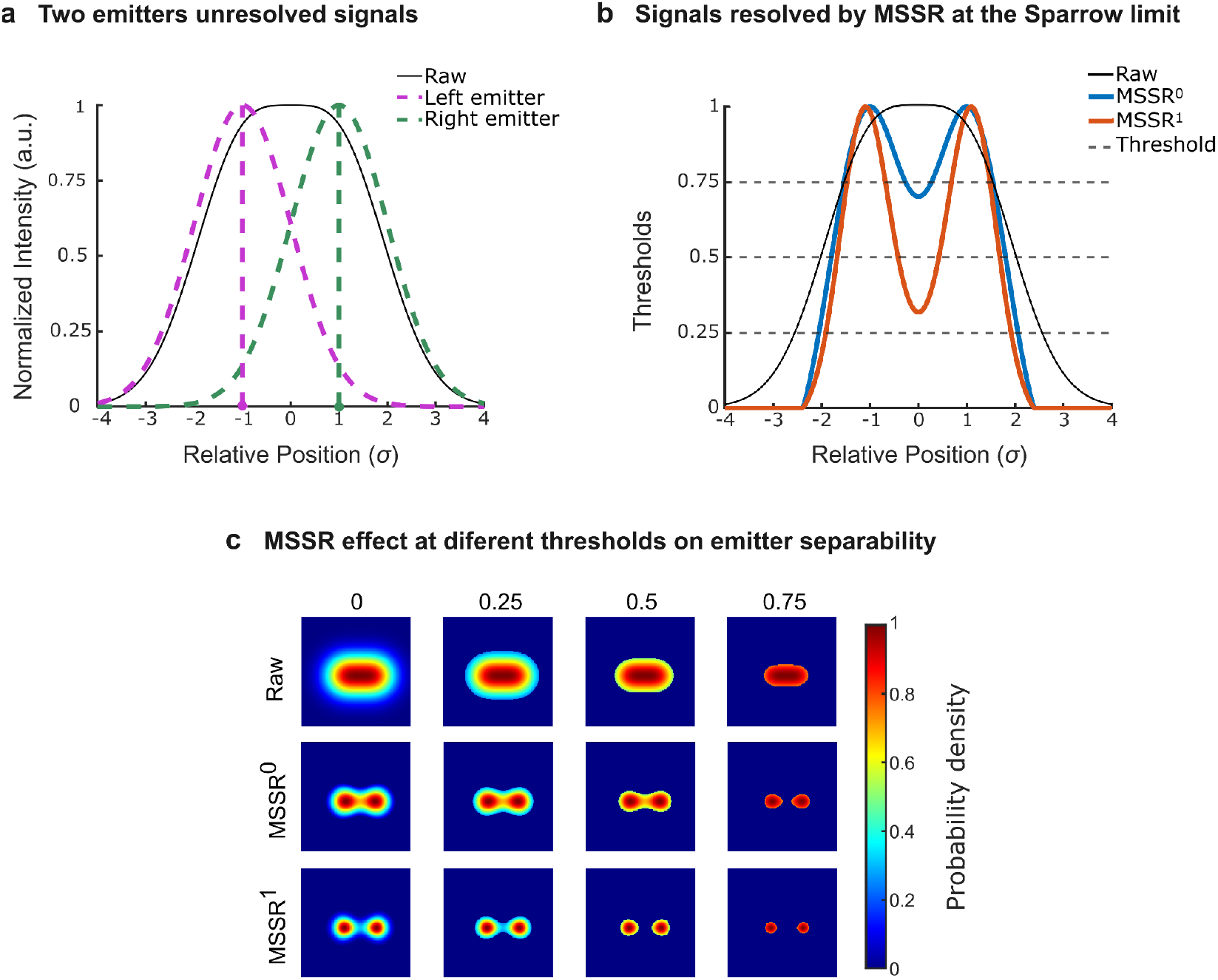
Resolution enhancement by MSSR at the Sparrow limit using a probability-like intermediate output (P_MSSR_). (a) Two Gaussian emitters separated by 2*σ* (Sparrow criterion) shown as individual contributions (magenta dashed, left emitter; green dashed, right emitter) and their summed signal (black). The summed profile is effectively unresolved at the midpoint under this separation. (b) One-dimensional profiles after MSSR processing, displayed as the intermediate bounded output used in this work P_MSSR_; computed prior to the final MSSR intensity remapping step). Raw (black), MSSR^0^ (blue), and MSSR^1^ (red) are shown. Horizontal dashed lines indicate representative threshold levels applied to P_MSSR_ to restrict support to high-probability regions. (c) Two-dimensional probability-density maps illustrating the combined effect of MSSR order (rows: Raw, MSSR^0^, MSSR^1^ and threshold level (columns) on emitter separability. Color bar indicates probability density (0–1). The horizontal axis in (a,b) is expressed in units of *σ*, the standard deviation of the Gaussian distributions used in panel a.

**Fig. 4.**
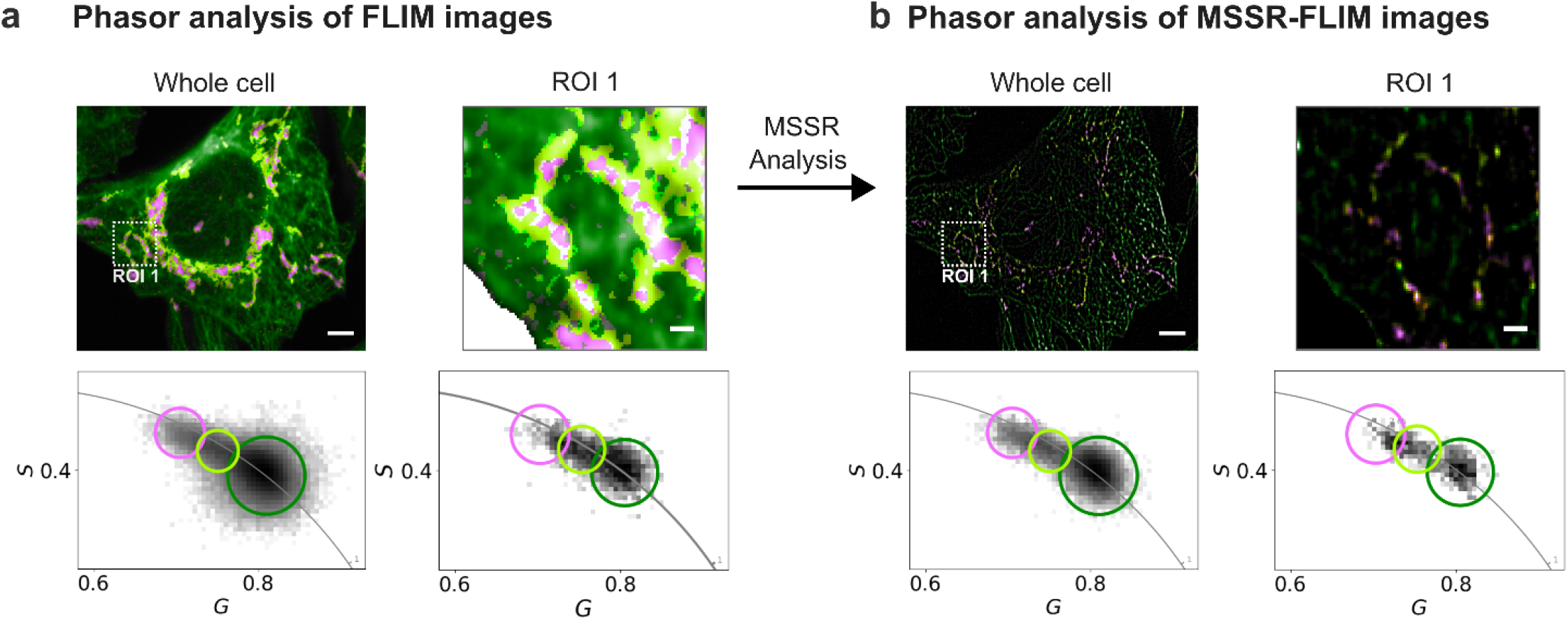
MSSR-derived probability masking reduces mixed-pixel signatures in phasor space without shifting endpoint lifetime clusters. (a) Phasor-based pseudo-color maps for the raw FLIM analysis of the whole cell and an inset ROI (ROI 1). The corresponding phasor plots (below) show endpoint-enriched lobes and an intermediate population. Circular cursors define the pseudo-color classes (green and magenta for endpoint-enriched classes; yellow for the intermediate/mixed class). (b) MSSR-masked analysis (“MSSR-FLIM”): MSSR is applied to an intensity projection to generate a bounded probability-like map 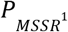, which is thresholded to form a binary mask that is applied to the phasor-coordinate images (G,S) computed from the unmodified time-resolved FLIM data. Endpoint loci remain stable while the mapped spatial support shows improved separability in ROI 1 and reduced intermediate-class pixels. Centroid displacement and the metrics associated with the compactness of pixel distributions in the phasor plots are reported in the main text and Methods. Scale bar in whole cell image: 3μm; scale bar in ROI: 500nm.

To test this prediction in the cellular specimen, MSSR was applied to an intensity projection of the FLIM acquisition to generate 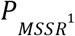, which was thresholded to produce a binary mask. The mask was then applied to the phasor-coordinate images (G,S) computed from the original time-resolved FLIM data, so that only spatially supported pixels contributed to the phasor plot and to the corresponding pseudo-color maps (Fig. 4b). If MSSR preprocessing had distorted decay kinetics, such distortion would manifest as a systematic displacement of the endpoint phasor clusters. Instead, the masked analysis preserves the endpoint loci while altering the spatial sampling of pixels that enter the analysis (compare Fig. 4a versus Fig. 4b), consistent with conservation of the underlying lifetime signatures. This invariance was quantified by computing phasor-cluster centroids before and after masking and measuring their displacement as a Euclidean distance in (G,S) space; centroid shifts were minimal (0.0066 for the whole cell and 0.0064 for ROI 1) and remained well below a conservative no-distortion bound of 0.05 phasor units selected to exceed fluctuations expected from photon noise and finite sampling (Fig. 4a–b; Methods). In parallel, the spatial maps show improved delineation of microtubule-like and mitochondria-like structures within the ROI, accompanied by an apparent reduction of pixels assigned to the intermediate mixed class (Fig. 4b, ROI 1). Consistent with reduced mixing, the phasor clusters obtained after masking exhibit increased compactness, which was quantified using an ellipse-area ratio defined as the cluster ellipse area after masking divided by the area before masking, which remained below unity for endpoint clusters (Methods). Because reduced phasor cluster area reflects diminished pooling of heterogeneous decays within single pixels, the combined behavior of minimal centroid displacement and reduced ellipse area supports the conclusion that MSSR-derived probability masking mitigates TB-associated mixing signatures while preserving the endpoint lifetime components of the specimen. In other words, it provides a principled way to reduce intermediate apparent lifetimes attributable to photon mixing.

### In-silico characterization of diffraction-induced Temporal Blending evidenced as phasor mixing

To systematically characterize TB, we developed an *in silico* model composed of two independent, non-interacting fluorophores. This simulation isolates and quantifies the contribution of point spread function (PSF) overlap to TB at controlled spatial separations (Fig. 5). The model is applicable to both time-domain FLIM (TD-FLIM) and frequency-domain FLIM (FD-FLIM) modalities.

**Fig. 5.**
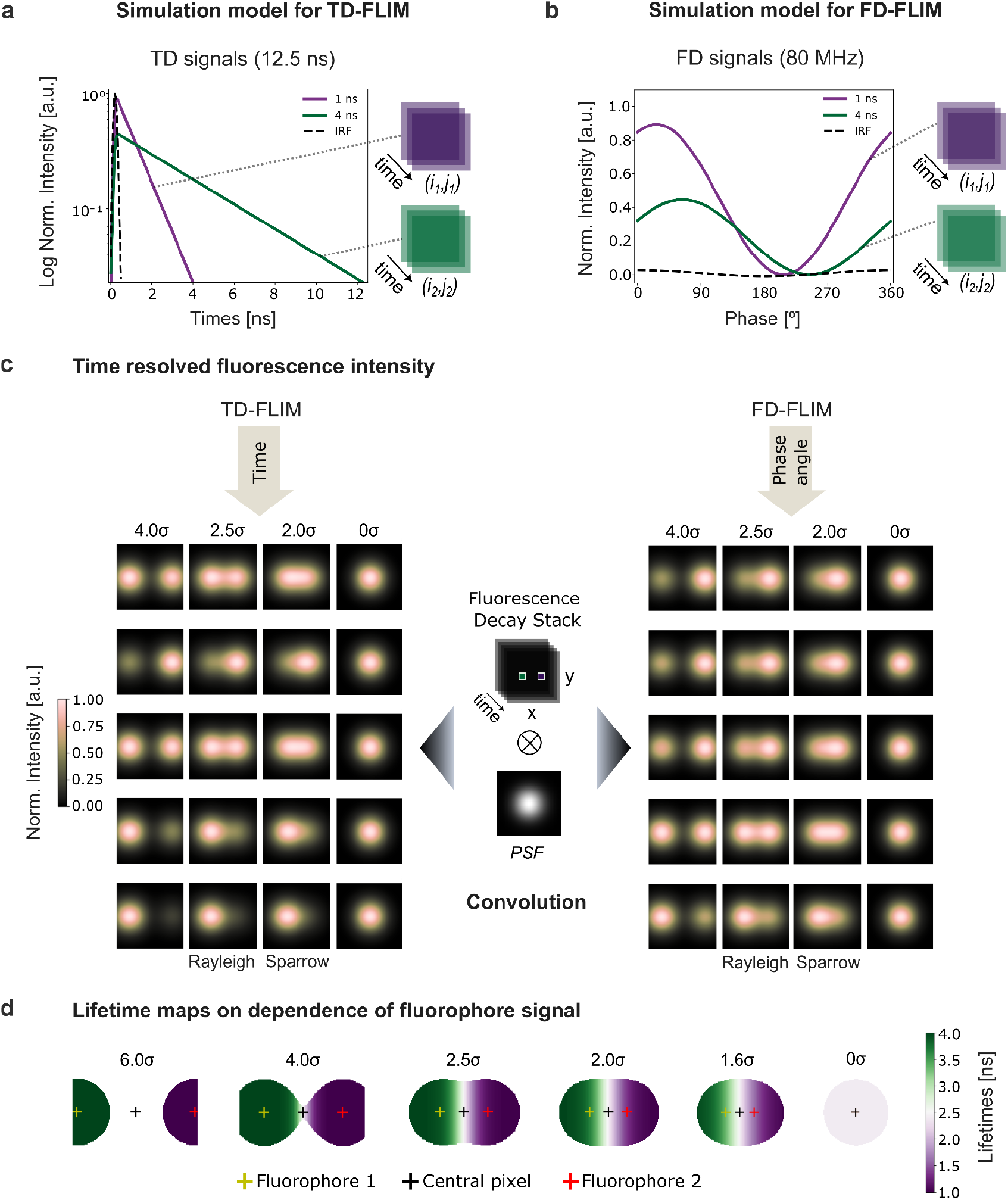
Simulation framework for characterizing Temporal Blending in FLIM. (a, b) Schematic representation of the *in silico* model for two independent, non-interacting fluorophores in TD-FLIM (a) and FD-FLIM (b). The simulated time-varying fluorescence signal of each fluorophore is stored in a single pixel at a defined position (*i*_*k*_, *j*_*k*_ ). (c) To account for diffraction-limited spatial blurring, both pixels are convolved with a Gaussian point spread function (PSF), yielding time-resolved image stacks; representative frames are shown for separations of 4.0σ, 2.5σ, 2.0σ, and 0σ. (d) After temporal analysis using the Fourier-based phasor workflow, pseudo-color lifetime maps are generated for separations from 6σ to 0σ, showing progressive emergence of intermediate modulation apparent lifetimes in the overlap region. Crosses indicate fluorophore positions (fluorophore 1, long-lived; fluorophore 2, short-lived) and the central reference pixel (empty unless the separation is 0σ).

Building on an initial frequency-domain approximation of signal mixing [53], we implemented a more general simulation framework that explicitly accounts for PSF-induced mixing prior to phasor extraction. This refinement avoids a coarse-grained representation of temporal behavior and enables the same simulation backbone to be used in both TD-FLIM and FD-FLIM regimes. Using PhasorPy [54], time-resolved fluorescence signals were generated for two fluorophores with distinct lifetimes of 1 ns and 4 ns. The temporal information was initially assigned to a single pixel for each fluorophore, namely fluorophore 1 (long-lived, 4 ns) at (*i*_1_, *j*_1_) and fluorophore 2 (short-lived, 1 ns) at (*i*_2_, *j*_2_) (Fig. 5a,b). A time-resolved image stack with dimensions (*x, y, t*) was then constructed, where each frame *t* represents a discrete time point within the simulated excitation cycle. To model spatial blurring, the fluorophore pixels were convolved with a two-dimensional Gaussian PSF (Fig. 5c).

Figure 5c illustrates the time-varying fluorescence intensities in both regimes and the corresponding blurred frames for representative separations (4.0σ, 2.5σ, 2.0σ, and 0σ). In TD-FLIM (left), intensity decreases monotonically across frames, while in FD-FLIM (right), the signal oscillates sinusoidally with the expected phase shift behavior. After temporal analysis of each simulated condition using the same Fourier-based phasor workflow described in the Methods and illustrated in Figure 6, pseudo-color lifetime maps were generated for six spatial separations ranging from 6σ to 0σ (Fig. 5d). When the fluorophores are well separated, pixels within each PSF report the ground-truth lifetimes (1 ns and 4 ns). As the fluorophores approach one another, PSF overlap increases and lifetime values blend in the overlap region, producing intermediate apparent lifetimes. In the case of complete spatial overlap (0σ), the phasor representation collapses toward a single amplitude-weighted mixed point, yielding a spatially uniform apparent lifetime in the derived map.

**Fig. 6.**
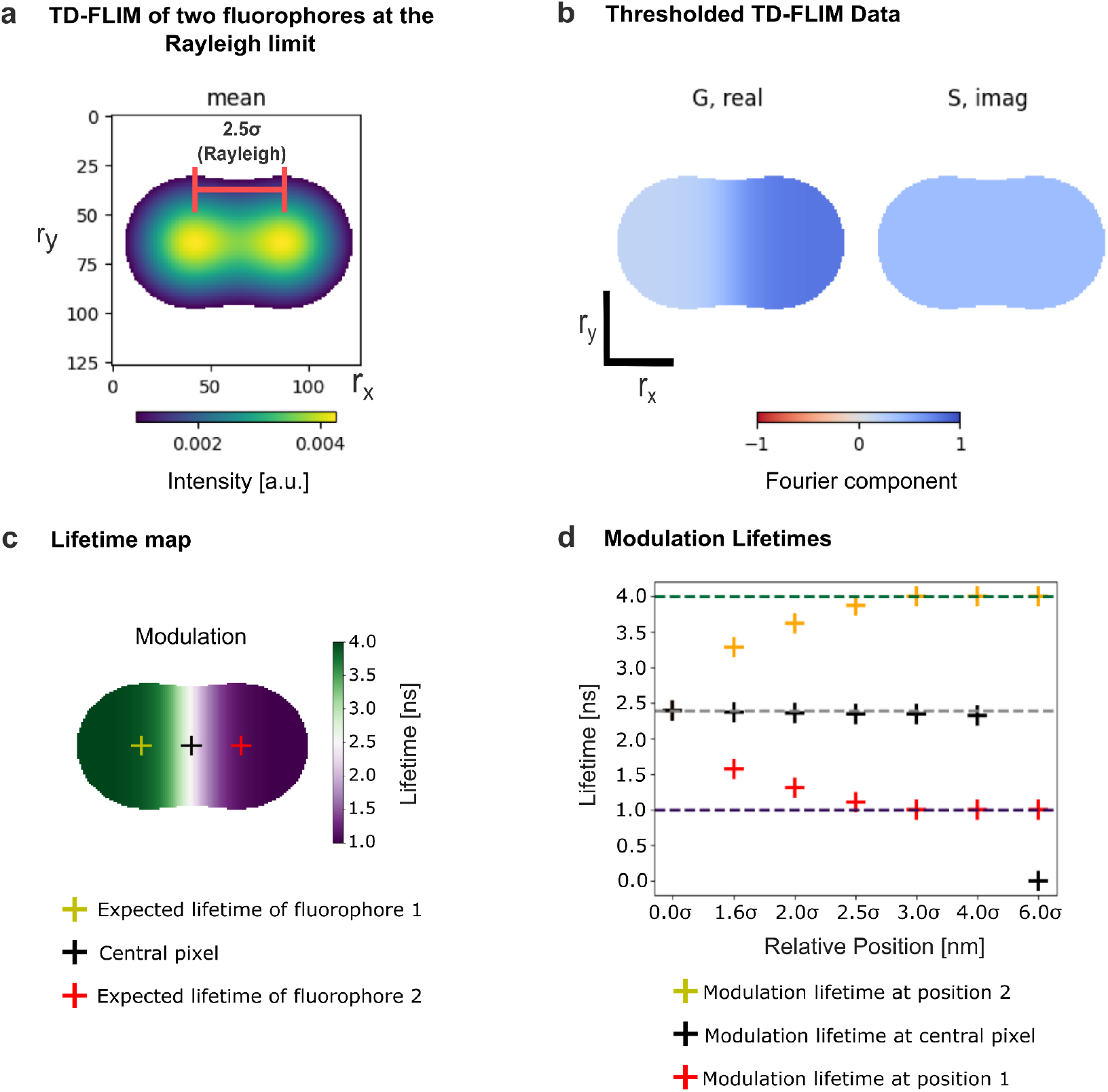
Temporal Blending in FLIM arises from PSF-induced spatial blurring. Panels (a–c) show simulated TD-FLIM images of two fluorophores positioned at 2.5σ. (a) Steady-state fluorescence intensity (mean from Fourier analysis) with intensity thresholding applied to suppress background. Axes represent pixel coordinates (*r*_*x*_, *r*_*y*_ ). (b) Phasor component maps: G (real) and S (imaginary). (c) Modulation lifetime map computed from Fourier components. (d) Modulation lifetimes sampled at the centers of each fluorophore position and at the central reference pixel, plotted as a function of inter-fluorophore separation (σ).

For the lifetime analysis, each simulated FLIM stack was processed using Fourier decomposition to transform temporal fluorescence signals into phasor space (Fig. 6). The time-invariant component corresponds to the steady-state fluorescence (mean intensity), illustrated for the case of two fluorophores positioned at 2.5σ (Fig. 6a). Phasor coordinates were computed using the Fourier-based implementation in PhasorPy [54]. Background contributions were suppressed by applying an intensity threshold to the mean intensity map (Fig. 6a).

Phasor component maps (Fig. 6b) revealed a smooth gradient in the G component between the two simulated fluorophores rather than a sharp boundary, consistent with PSF-driven mixing of temporally distinct signals. The corresponding modulation lifetime map (Fig. 6c) further confirms this effect through the emergence of intermediate apparent lifetimes in the overlap region. Notably, the phase lifetime map displays a shift in the transition zone toward the long-lifetime fluorophore (Supplementary figure S2), suggesting a similar effect to the Frequency Modulation Capture Effect (FMCE) described by Yuan, X. et al. [55]. Finally, modulation lifetime values sampled at the centers of the fluorophore positions and at the central reference pixel were quantified as a function of separation (in units of σ) (Fig. 6d). A clear dependence on inter-fluorophore separation was observed, highlighting the role of PSF overlap in driving TB in FLIM measurements. The same framework was used to study TB in the FD-FLIM simulations (Supplementary figure S3).

TB remains visible in pseudo-color lifetime maps even at an inter-fluorophore distance of 4σ (Fig. 7). The case at 1.6σ is particularly informative, because this separation has been reported as a practical MSSR resolution limit in [51]. As the separation decreases from well-resolved to fully overlapped, the phasor cluster broadens and shifts toward intermediate G values, consistent with an amplitude-weighted phasor mixture produced by PSF overlap and photon pooling, which yields apparent intermediate lifetimes in the derived maps. In the fully overlapped case (0σ), the phasor distribution collapses to a single point corresponding to the mixture of both components. At the opposite extreme (6σ), only one pixel is sampled for each lifetime, as the modeled signal arises solely from the two discrete fluorophores with no additional contributions.

**Fig. 7.**
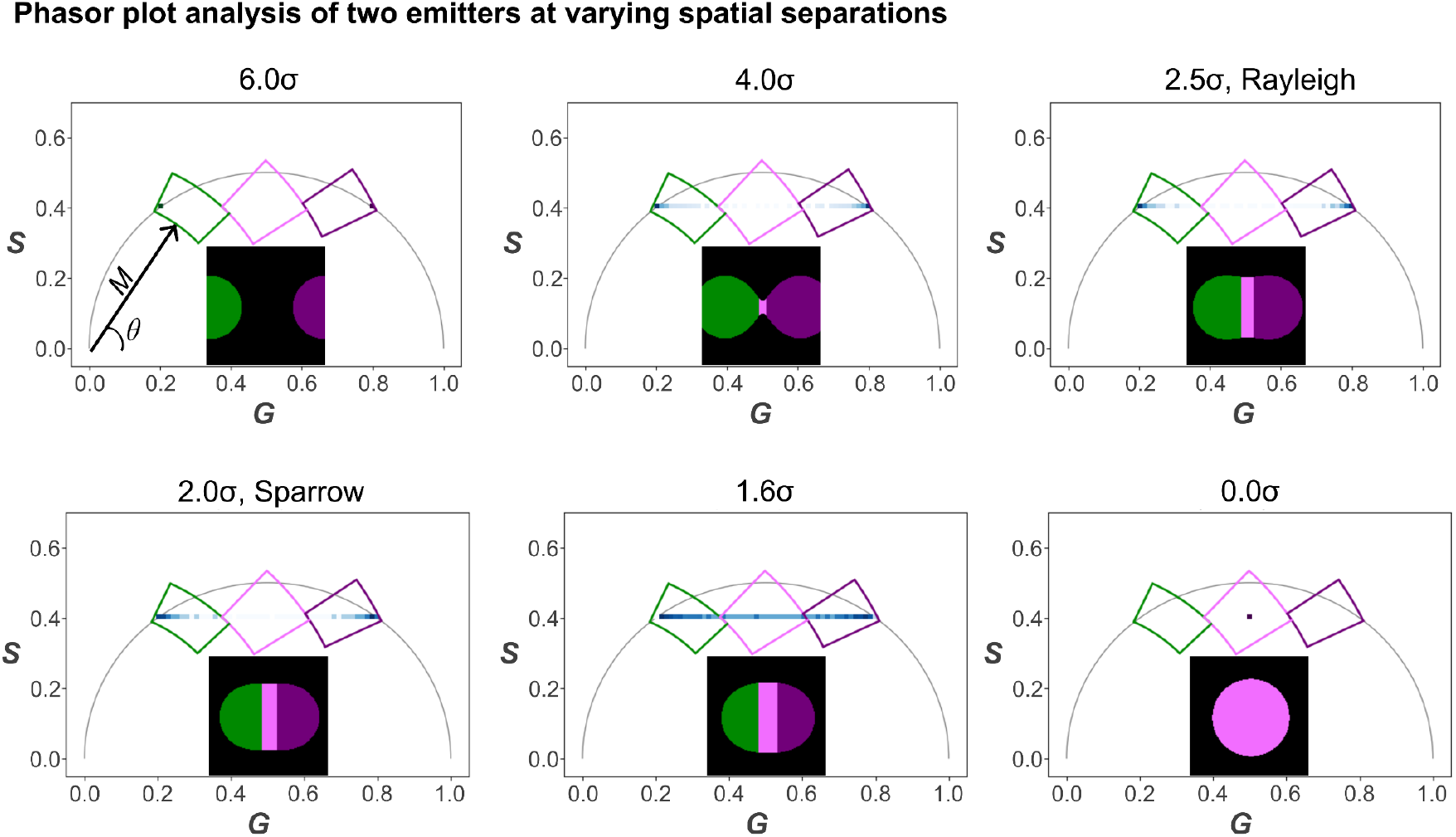
Phasor analysis across inter-fluorophore separations reveals progressive broadening and migration of mixed-pixel signatures. Each panel shows the phasor distribution (G, S) for a given separation (in units of σ) with the universal semicircle. Rectangle-polar cursor selections illustrate how phasor-space regions map, via reciprocity, to spatially distinct regions in the corresponding pseudo-color lifetime maps (insets). Increased PSF overlap produces intermediate phasor positions consistent with temporally mixed pixels.

Following the approach described above, MSSR was applied to reduce TB and enhance spatial resolution for the challenging case of two emitters positioned at the Sparrow diffraction limit (2σ), shown in Figure 8. As illustrated in Figure 8a, the raw intensity image exhibits substantial PSF-induced signal overlap, which manifests in phasor space as an extended distribution characteristic of mixed pixels (Fig. 8a, right). Cursor selection was used to isolate pixels corresponding to monoexponential decays, operationally defined as phasor components lying along the universal semicircle. The associated modulation lifetime map (Fig. 8a, middle) reveals a spatially continuous intermediate lifetime region between the two emitters, reflecting temporal blending arising from spatial averaging. MSSR was then applied as a spatial probability mask derived from the fluorescence intensity distribution, using percentile-based thresholds (25th, 50th, and 75th) to progressively isolate regions with a high likelihood of originating from individual emitters; each threshold mask was applied prior to the phasor transformation (Supplementary Fig. S4). Figure 8b shows the result obtained using the 75th-percentile threshold. Under this Sparrow-limit condition, the MSSR markedly improves spatial separability, as evidenced in the modulation lifetime map by the collapse of the intermediate lifetime region and the recovery of spatially confined lifetime domains associated with each emitter. In phasor space, this corresponds to a pronounced compaction of the phasor distribution. Quantitatively, this reduction is captured by a strong decrease in phasor cluster area, expressed as an ellipse area ratio (*A*_*after*_/*A*_*before*_) of 1.15 × 10^-9^, together with a minimal centroid displacement of 0.0045 in (G, S) units, indicating effective suppression of TB while preserving the intrinsic decay signatures.

**Fig. 8.**
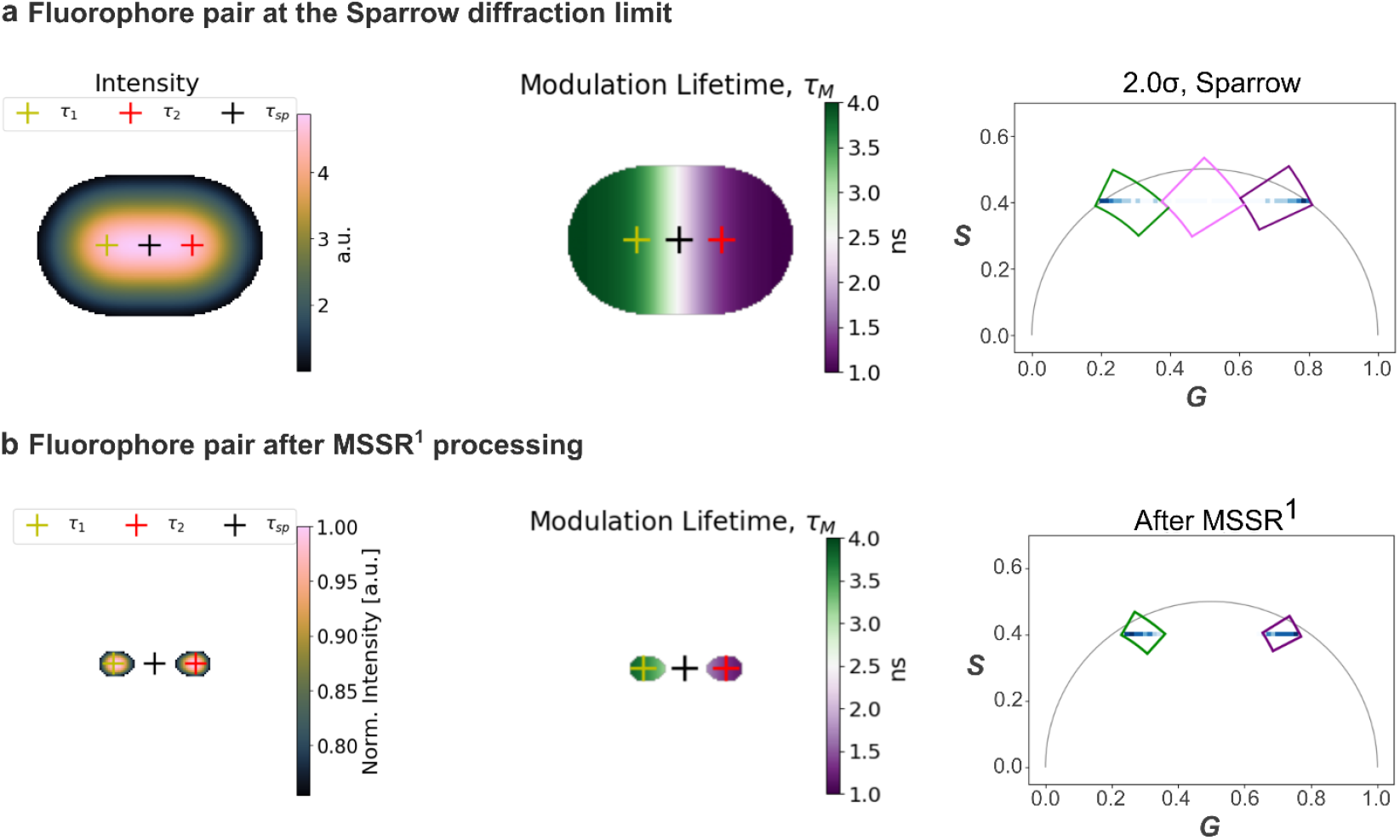
Mitigating Temporal Blending using Mean-Shift Super-Resolution: a computational approach. Two independent, non-interacting simulated fluorophores are positioned at the Sparrow limit (2.0σ; Fig. 3). (a) Raw condition: intensity profile (left), modulation lifetime map (middle), and phasor plot (right) showing an extended distribution characteristic of mixed pixels; cursor selection highlights monoexponential components along the universal semicircle. (b) MSSR masking applied to the dataset in (a) prior to phasor transformation using the 75th-percentile threshold (see Fig. S4 for threshold comparison). The MSSR-masked analysis increases spatial separation and compacts the phasor distribution, indicating reduced Temporal Blending while preserving the component decay signatures; compaction is quantified by the ellipse-area ratio and centroid displacement in (G, S) units.

### Mean-Shift Super-Resolution addresses spatio-temporal resolution in three-component FLIM images

In the preceding sections, we demonstrated that masking FLIM images with MSSR-enhanced intensity maps mitigates temporal blending arising from the spatial overlap of two independent fluorophore PSFs. Here, we extend this approach to a more complex three-component system and show that the same strategy reduces lifetime mixing originating from spatial overlap of three distinct fluorescent species.

Figure 9 presents a three-component TD-FLIM image of a single U2OS cell labeled with DAPI (blue), AF555 (green), and AF532 (yellow), staining the nucleus, microtubules, and mitochondria, respectively (same sample as in Figures 1, 2, and 4, but acquired using a different instrument and imaging configuration, see methods). Figure 9a shows the diffraction-limited FLIM image of the whole cell prior to MSSR masking. The white square delineates a region of interest (ROI) containing pixels where the three labeled subcellular structures spatially overlap. The corresponding whole-cell phasor plot is presented for reference and exhibits a characteristic triangular distribution, indicative of a three-component lifetime system. Each vertex of the triangle corresponds to the intrinsic lifetime of one fluorophore: the left vertex to AF532 (longest lifetime), the central vertex to AF555, and the right vertex to DAPI (shortest lifetime). The pseudo-color encoding of the lifetime image reflects the cursor-based segmentation defined in the phasor plot.

**Fig. 9.**
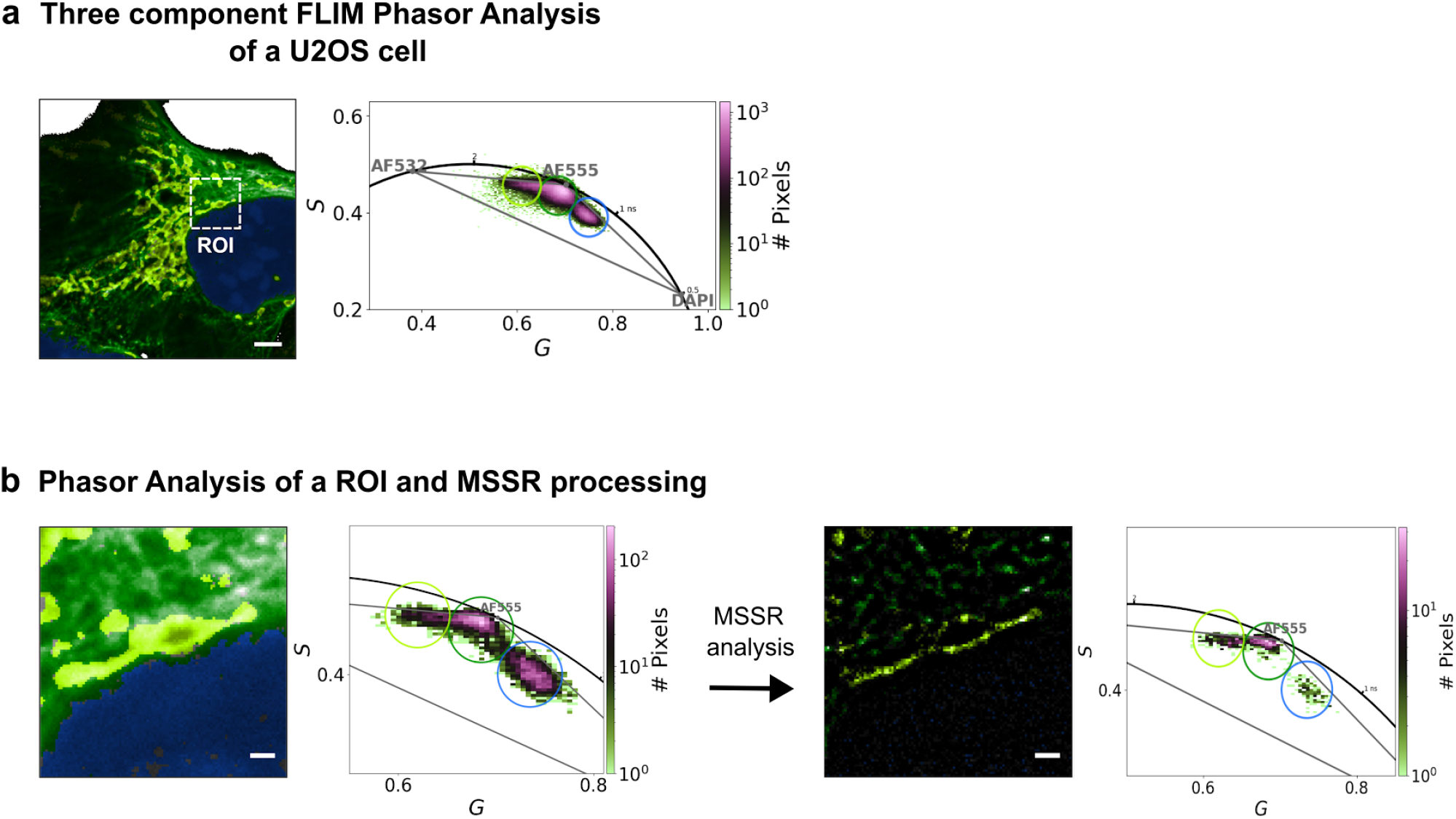
Mitigating temporal blending using Mean-Shift Super-Resolution (MSSR) on a three-component FLIM image. (a) Three-component FLIM phasor analysis of a single U2OS cell labeled with DAPI (blue, nucleus), AF555 (green, microtubules), and AF532 (yellow, mitochondria). Left: diffraction-limited pseudo-color FLIM image of the whole cell with the ROI indicated (white square). Right: whole-cell phasor plot showing a triangular distribution characteristic of a three-component lifetime system, with vertices corresponding to AF532 (left; longest lifetime), AF555 (center), and DAPI (right; shortest lifetime). Pseudo-color encoding reflects the cursor-based segmentation defined in the phasor plot. (b) ROI phasor analysis and MSSR processing. Left: diffraction-limited ROI image and the corresponding ROI phasor distribution (pre-MSSR). Middle: MSSR analysis step. Right: MSSR-masked ROI image using the 8% intensity threshold and the corresponding ROI phasor distribution computed after masking (post-MSSR). MSSR reduces mixed pixels while preserving phasor cluster positions. Scale bar in whole cell: 4 μm; scale bar in ROI: 800nm.

Figure 9b presents the ROI-level analysis used to evaluate MSSR masking. The left side of Figure 9b shows the diffraction-limited ROI (image) and its corresponding ROI phasor distribution (pre-MSSR), where mixed pixels populate the interior and edges of the triangular manifold expected for a three-component system. The right side of Figure 9b shows the same ROI after MSSR masking using an 8% intensity threshold, together with the corresponding ROI phasor distribution computed after masking (post-MSSR). Under these conditions, MSSR reduces spatial overlap between fluorophore signals, thereby enhancing spatial separation in the ROI FLIM image. The centroids of the three phasor clusters remain stable, as confirmed by the centroid displacement metric (0.034), which remains below the predefined tolerance of 0.05 and therefore indicates no significant alteration of intrinsic fluorescence decay dynamics. Consistently, a modest reduction in the phasor ellipse area ratio (0.96) is observed after MSSR processing, indicating a small global decrease in lifetime mixing within the ROI phasor distribution. Notably, the segment connecting the DAPI and AF555 phasor populations shows a pronounced depletion of intermediate pixels, yielding two more distinct clusters and indicating suppression of temporal blending along this axis.

These findings establish that Mean-Shift Super-Resolution enables spatial refinement of FLIM images without perturbing the underlying lifetime information, even in complex, multi-component systems. The persistence of stable phasor centroids alongside reduced lifetime mixing supports a spatial—rather than temporal—origin of Temporal Blending.

## DISCUSSION

Temporal blending in FLIM has been approached from multiple theoretical and computational perspectives, yet a unified interpretation that cleanly separates spatial from temporal contributions has remained incomplete. Here, we frame TB primarily as a PSF-driven mixing effect and show that application of MSSR-generated masks reduces this mixing while preserving intrinsic lifetime information, as evidenced by phasor-cluster centroid stability together with its reduced area.

Early deconvolution-based strategies, such as the Tikhonov–Miller reconstruction introduced by Squire and Bastiaens [18], addressed spatial blurring by applying regularization independently to each Fourier component (mean, real, and imaginary frames) to deblur FLIM images by minimizing a Tikhonov functional [50]. While effective at sharpening images, regularization performed directly on time-varying components can differentially reshape harmonics under low signal-to-noise ratio and/or elevated background, which may distort phasor geometry and bias lifetime interpretation by mimicking genuine heterogeneity.

A complementary analytical framework was introduced by Yuan et al. [55] through the Frequency Modulation Capture Effect (FMCE), which describes how, in FD-FLIM, multi-emitter measurements can become dominated by the strongest emitter. In that model, amplitude-dependent capture biases the measured lifetime toward the brightest fluorophore, producing smoothly varying lifetime distributions and spatial mislocalization artifacts. To mitigate this effect, Yuan et al. proposed a hybrid strategy in which spatial resolution is enhanced using SRRF on an intensity image and subsequently combined with the lifetime map. Although powerful, this approach typically relies on multi-frame reconstruction and temporal interpolation steps when applied to FLIM workflows, and it can be computationally demanding because SRRF benefits from large frame counts for optimal performance. Importantly, amplitude capture (FMCE) and PSF-weighted spatial mixing are not mutually exclusive; both can co-occur in experimental data. The present study isolates the geometric–optical contribution by using *in silico* conditions in which fluorophores are non-interacting and have monoexponential fluorescence decay, so that apparent lifetime heterogeneity arises from spatial convolution and not from environmental parameters in the vicinity of the fluorophore.

Furthermore, the present work demonstrates that applying MSSR exclusively to the intensity (DC) image prior to phasor transformation markedly reduces TB while preserving intrinsic fluorescence lifetimes. MSSR generates an intensity-derived spatial discrimination mask that emphasizes emission maxima and suppresses low-intensity neighborhoods where PSF overlap contributes disproportionately to mixing. Phasor coordinates are then computed on the masked pixel set, without temporal interpolation, resampling, or frequency-domain manipulation of the decay. Consequently, lifetime estimation remains tied to the measured temporal information, while the degree of spatially induced mixing is reduced upstream of the lifetime transform.

Crucially, phasor analysis indicates that the centroids of phasor clusters remain practically invariant before and after MSSR masking, supporting preservation of intrinsic lifetime information. Simultaneously, the marked reduction in phasor-cluster dispersion and improved separation between components demonstrate that MSSR suppresses overlap-driven mixing without introducing synthetic decay components. This behavior distinguishes MSSR from approaches that act directly on time-varying frames or implicitly modify temporal content through reconstruction.

The *in silico* characterization further reveals that TB persists even when fluorophores are spatially resolvable by conventional diffraction-based criteria. Significant phasor mixing remains detectable at emitter separations as large as 4σ, intensifies near the 2σ Sparrow limit, and becomes maximal at intermediate separations around 1.6σ for the simulated conditions. At complete overlap (0σ), the phasor distribution collapses toward the intensity-weighted linearly-combined phasor of the overlapping lifetimes. These effects emerge despite the presence of strictly non-interacting emitters, having monoexponential relaxation dynamics demonstrating that apparent lifetime heterogeneity can arise purely from PSF-mediated spatial blending. The precise separation thresholds will depend on SNR, background signals, relative emitter brightness, sampling, and the PSF model.

Based on these results, we propose a mechanistic interpretation in which TB arises from PSF-weighted spatial averaging imposed by the imaging system prior to lifetime estimation (a formal treatment is provided in the Supporting Information). When neighboring emitters fall within each other’s diffraction volumes, their photon contributions are aggregated into single measurement pixels, yielding an apparent decay that reflects a spatially weighted superposition rather than a true molecular lifetime distribution at a single location. The persistence of phasor mixing at separations as large as 4σ supports this geometric–optical origin and motivates mitigation strategies that reduce effective spatial averaging before the lifetime transform.

The experimental results corroborate this mechanism across two- and three-component datasets at whole-cell and ROI scales. MSSR consistently reduced phasor-cluster area and preserved its centroid positions. In the most challenging three-component system, MSSR improved the separation of AF555 and DAPI populations, reducing extensive mixing observed in diffraction-limited FLIM and yielding more symmetric, well-separated phasor distributions.

It is instructive to contrast this approach with super-resolution strategies based on higher-order correlations, such as antibunching-based microscopy proposed by Schwartz and Oron [56]. In that framework, spatial resolution is enhanced by exploiting the nonclassical photon statistics of fluorescence emission, effectively narrowing the PSF through higher-order intensity correlations. While conceptually distinct, both approaches highlight the central role of spatial statistics in overcoming diffraction-induced averaging. However, antibunching-based methods require photon-number-resolving detection and high photon fluxes, whereas MSSR operates entirely within the classical intensity domain and can be readily applied to standard FLIM datasets without specialized hardware or acquisition schemes.

Together, these results demonstrate that MSSR alleviates the geometric-optical origin of Temporal Blending while preserving the physical decay properties of fluorophores. By decoupling spatial resolution enhancement from temporal manipulation, MSSR resolves the long-standing spatial–temporal trade-off in FLIM. The method therefore provides a computationally simple, modality-agnostic strategy that complements phasor based lifetime analysis and expands the practical limits of super-resolved FLIM imaging.

## CONCLUSIONS

This study identifies and addresses a fundamental limitation in fluorescence lifetime imaging microscopy (FLIM), namely the intrinsic coupling between spatial resolution and temporal accuracy caused by Temporal Blending. Through combined *in silico* modeling and experimental validation, we demonstrate that this effect is driven by optical PSF-mediated spatial averaging, which produces apparent lifetime mixing even for strictly monoexponential emitters.

By applying Mean-Shift Super-Resolution (MSSR) as a spatial pre-processing step prior to phasor transformation, we show that TB can be substantially mitigated without introducing lifetime artifacts. MSSR acts as a probabilistic spatial filter operating exclusively on intensity information, thereby preserving intrinsic fluorescence decay kinetics. Phasor analysis confirms this preservation through practically invariant centroid positions, and reduced phasor cluster spread corresponding to reduced temporal mixing. In contrast to iterative deconvolution or temporally interpolative super-resolution approaches, MSSR enhances spatial resolution without distorting phasor geometry or decay information. Another strength of MSSR is its capability of enhancing spatial resolution within one frame of microscopy images, thus lowering significantly the computational resources required to process the data.

Beyond its practical utility, this work proposes a mechanistic interpretation in which TB arises predominantly from PSF-weighted spatial convolution. While this model is strongly supported by the present simulations and experimental observations, further validation across diverse fluorophore systems, imaging modalities, and noise regimes will be necessary to fully establish its generality. Nevertheless, this interpretation provides a physically grounded framework for understanding spatial-temporal coupling in FLIM and suggests new pathways for redefining resolution limits in lifetime imaging.

Overall, the MSSR-FLIM framework offers a conceptually robust and computationally efficient strategy for improving spatial resolution while preserving temporal fidelity. These results also highlight the phasor approach as an effective tool to diagnose and quantify the influence of spatial filtering on lifetime data. The methodology is immediately applicable to live-cell FLIM, super-resolution workflows, and lifetime unmixing, offering a generalizable strategy to overcome spatial–temporal trade-offs in modern fluorescence microscopy.

## Materials and Methods

All computational analyses were implemented as a modular, notebook-based workflow in the public MSSR-FLIM-2025 repository hosted on GitHub. The repository code/ directory contains multiple notebooks encoding distinct processing stages, including instrument export handling, phasor computation and visualization, cursor-based segmentation, MSSR-based spatial masking, synthetic simulations, and downstream quantification. The Materials and Methods section is structured to mirror this modular organization, such that each methodological block is self-contained and can be extended in later subsections as additional analyses are introduced. Repository: https://github.com/MarioGoG/MSSR-FLIM-2025

### Dataset, specimen preparation, and labeling strategy

Primary experimental analyses were performed on time-domain confocal FLIM data acquired from a standardized multicolor FLIM slide containing fixed human osteosarcoma U2OS cells (GATTA-Cells 3C FLIM; GATTAquant). The slide comprises three spectrally distinct labels: nuclear DNA labeled with DAPI, mitochondria labeled by TOM20 immunolabeling with Alexa Fluor 532, and microtubules labeled by α-tubulin immunolabeling with Alexa Fluor 555. Samples were embedded in ProLong Diamond antifade medium and sealed.

The immunolabeling strategy immobilizes fluorophores on distinct macromolecular targets (TOM20-positive mitochondrial membranes versus α-tubulin–positive microtubules). For the Alexa Fluor 532 and Alexa Fluor 555 channels, this supports a working model of two spatially interlaced yet non-coupled lifetime species at the scale of labeled structures. The same physical slide was used for two-color FLIM (Alexa Fluor 532 and Alexa Fluor 555) and, when required, three-color FLIM including the DAPI channel. Channels were processed independently using an identical phasor-analysis formalism.

### FLIM image acquisition

Time-resolved confocal FLIM images were acquired on a custom microscope platform (iRT-FLIM-001; Intek Scientific) described previously. Excitation was provided by a 515 nm pulsed laser (Prima; PicoQuant) operated at a 25 MHz repetition rate. Images were collected using a 60× objective (UPLXAPO60; Olympus), a 0.5 Airy unit pinhole, and a pixel size of 54 nm. Emission was filtered to 550–614 nm using a bandpass filter (FF01-582/64-25; Semrock) and detected with a photomultiplier tube (H10720-01; Hamamatsu Photonics). For each pixel, the acquisition yielded a time-domain fluorescence decay sampled over a fixed nanosecond-scale temporal window. A modulation of 50 MHz (2nd harmonic) was used for the calculation of G and S phasor coordinates. More details on the experimental set up can be found on [57].

Selected datasets were additionally acquired on a Leica TCS SP8 STED 3X platform equipped with a FALCON-FLIM module (photon-counting hybrid detectors, galvanometric scanning, and a 63× high-numerical-aperture oil-immersion objective). The acquisition frequency for this system was 78 MHz. These datasets were analyzed with the same conceptual workflow (phasor-based segmentation with optional MSSR-derived spatial masking); instrument-specific acquisition parameters were recorded per experiment and are described in the methods section “Three-component TD-FLIM phasor analysis and MSSR masking”.

### Phasor computation, exported products, and data import

Phasor analysis followed the standard FLIM phasor formalism and Weber identities. Time-domain decays were converted to per-pixel phasor coordinates (g,s) and mean intensity (DC component), yielding spatial matrices for mean intensity and the phasor real and imaginary components. Dual modulation frequencies (50 MHz and 100 MHz) were used for phasor computation, implemented via cosine and sine transforms and accelerated on CUDA-enabled GPUs when available (LifetimeXplorer v1.10.9). For downstream processing in Python, mean intensity images (projections across the time-gated stack) were imported from TIFF (e.g., *_Mean.tif) using tifffile, and phasor coordinate matrices were imported from CSV (e.g., *_Real.csv, *_Imag.csv) using pandas.

For the Leica system, phasor analysis was performed using the manufacturer’s built-in software with a modulation frequency of 80 MHz. The raw time-correlated single-photon counting (TCSPC) data were exported in PicoQuant’s proprietary PTU file format. The results of the phasor analysis were exported as a Leica Image File (.lif), a proprietary Leica file format containing the real (G) and imaginary (S) phasor coordinates.

### Frequency conventions, harmonic support, and calibration

Downstream phasor operations used explicit frequency parameterization following PhasorPy conventions. A representative fundamental frequency was specified as frequency = 0.05 in GHz units, corresponding to *f* = 50∼*MHz* with ω = 2π*f*. Harmonic support was enabled by specifying a harmonic index (e.g., harmonic = 1 for the fundamental). When reference-based calibration was required, calibration utilities (e.g., phasor_calibrate) were applied using a reference lifetime-derived phasor and a measured reference phasor to correct systematic offsets.

### Intensity thresholding and spatial stabilization of phasor maps

To suppress noise-driven phasor scatter, pixels with insufficient photon counts were excluded before visualization, segmentation, and lifetime extraction. Thresholding was applied to the mean intensity image to generate a binary support mask that removed background and low-count regions. Unless otherwise stated, (g,s) and derived lifetime quantities were computed from the original time-resolved FLIM data; masking was applied during pixel selection, visualization, and class-restricted statistics. Representative preprocessing parameters included a minimum mean-intensity threshold (mean_min = 7) and optional Gaussian smoothing of mean intensity (scipy.ndimage.gaussian_filter, sigma = 1 pixel). To reduce salt-and-pepper noise in the phasor coordinate maps while preserving structural boundaries, repeated median filtering with intensity weighting was applied:

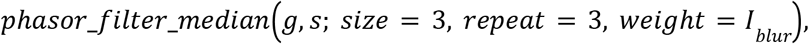

corresponding to a 3 × 3 spatial median filter iterated three times and weighted by the smoothed mean intensity.

### Phasor visualization and reciprocity mapping

Phasor distributions were rendered as density clouds in (*G, S*) space with a universal semicircle overlay for monoexponential decays. Monoexponential decays map to the semicircle, whereas multiexponential mixtures fall inside the semicircle due to amplitude-weighted linear combination behavior in phasor space. Operational reciprocity mapping was used bidirectionally: pixels selected by a phasor-space region were mapped back to image space to generate pseudo-color masks, and the inverse mapping was used to validate that spatially coherent structures corresponded to coherent phasor regions. Representative visualization settings included 2D histogram rendering with bins = 150 and axis limits tuned to the informative region (representative values: *g* ≈ 0. 55 − 1. 00, *s* ≈ 0. 00 − 0. 60).

### Cursor-based segmentation and pseudo-color classification

Segmentation in phasor space used circular cursors. For a cursor *k* centered at (*g*_*k*_, *s*_*k*_ ) with radius *r*_*k*_, pixel membership was defined as:

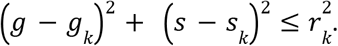

Two-component segmentation placed cursors at the two endpoint lobes of the phasor distribution to enrich for Alexa Fluor 555–dominant pixels (microtubules) and Alexa Fluor 532–dominant pixels (mitochondria). For three-class temporal blurring (TB) analysis, a third cursor was positioned over the intermediate region connecting endpoints to capture mixed pixels consistent with amplitude-weighted linear combinations. Representative full-field endpoint cursor centers were near (*G, S*) ≈ (0. 70, 0. 42) and (0. 82, 0. 40), with radii on the order of 0.07. For ROI-focused analyses, cursor centers and radii were refined to local phasor-density maxima, and numerical cursor sets were recorded explicitly for reproducibility. Cursor geometries and placements were held constant within unmasked versus masked comparisons to isolate the impact of spatial masking on phasor dispersion and class occupancy. Pseudo-color overlays were rendered by alpha blending categorical labels onto a grayscale mean-intensity background derived from the thresholded mean image. Spatial scaling used 54 nm per pixel for scale bars and overlays.

### ROI-based local subanalyses

To interrogate local interfaces between mitochondria and microtubules and to compare against regions dominated by a single structure, fixed ROIs were defined by array slicing of intensity and phasor matrices and processed with the same pipeline (thresholding, optional median filtering, phasor visualization, cursoring, pseudo-color mapping, and histogram-based quantification). Representative ROI definitions included ROI1, ROI2, …, ROIn.

### Apparent lifetime extraction from phasor coordinates

Apparent lifetimes were computed per-pixel (*G, S*) at modulation frequency *f* with ω = 2π*f*. Phase and modulation were computed as:

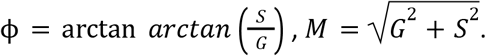

Phase lifetime (τ_ϕ_) and modulation lifetime (τ_*M*_) were computed via Weber identities:

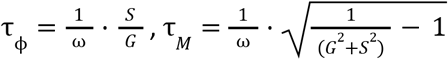

For monoexponential decays on the universal semicircle, τ_*M*_ is equivalent to τ_ϕ_.

Lifetime extraction was performed after intensity thresholding (and after median filtering when enabled). Downstream summaries emphasized modulation lifetime distributions.

### Lifetime histograms and Gaussian-mixture fitting

Lifetime distributions were summarized as histograms of modulation lifetimes τ_*M*_ over endpoint-enriched pixels or over endpoint plus intermediate TB pixels, depending on analysis objective. Two fitting regimes were used: (i) endpoint-only analyses with representative settings bins ≈ 95 and range ≈ 0. 6 − 2. 6, fitted with a two-Gaussian mixture; and (ii) endpoint plus intermediate analyses with representative settings bins ≈ 200 and range ≈ 0. 6 − 2. 6, fitted with a three-Gaussian mixture. Gaussian mixtures were fitted using scipy.optimize.curve_fit with parametric components:

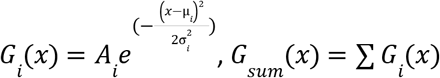

with physically plausible bounds on component means and widths. Fit initializations were selected from visually apparent histogram modes and recorded per ROI.

### Quantification of phasor preservation and dispersion changes after MSSR masking

Two complementary metrics were used to evaluate whether MSSR-based masking altered lifetime signatures or primarily reduced dispersion.

Centroid displacement (phasor stability) was computed for each endpoint cluster as the Euclidean distance between (*G, S* ) centroids before and after masking. To compute the phasor cloud centroids we used:

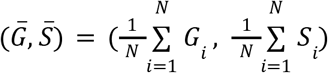

So that the Centroid displacement was calculated as:

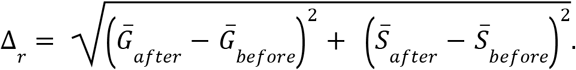

Displacements were interpreted relative to a conservative no-distortion bound of 0.05 phasor units, selected to exceed expected fluctuations from photon noise and finite sampling.

Cluster compactness (dispersion reduction) was summarized using confidence-ellipse area estimates. Compactness was quantified as an ellipse-area ratio:

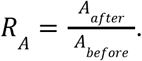

The ellipses areas were computed with the following equations, considering the estimation of an area with 95% confidence interval:

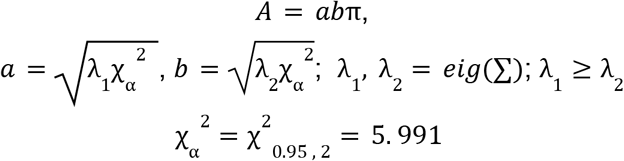

Values below unity indicate reduced dispersion. This metric was evaluated for whole-cell distributions and for ROIs to quantify local suppression of dispersion under stable endpoint coordinates. Together, these metrics distinguish (i) shifts in lifetime-coordinate endpoints, indicative of altered decay signatures, from (ii) reduced dispersion at stable endpoints, consistent with mitigated mixing while conserving endpoint lifetimes.

### Mean-Shift Super-Resolution as a masking strategy for spatial separability

Mean-Shift Super-Resolution (MSSR) was used as an intensity-domain spatial separability enhancer to generate spatial masks while preserving time-resolved decay information. In experimental FLIM analyses, MSSR was applied to the intensity projection of each acquisition (mean intensity image). MSSR follows local intensity gradients (mean-shift procedure) to concentrate probability density toward local maxima, thereby reducing effective point-spread-function overlap and improving separability of nearby structures without direct manipulation of the time-resolved decay.

For masking, the workflow used the intermediate MSSR output obtained prior to the final MSSR intensity remapping step, yielding a bounded matrix in the range [0,1], denoted 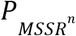. This intermediate output was treated as a probability-like spatial support map suitable for threshold-based retention of high-support pixels while suppressing low-support regions associated with background and overlap. Unless otherwise stated, masks were derived from the first iterative output 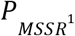 (corresponding to MSSR^1^ in the accompanying scripts). Binary masks were constructed by thresholding 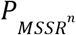 at selected levels. Threshold selection was explored using quartile-like cutoffs (0.25, 0.50, 0.75) in simulations and then transferred to experimental data using thresholds that retained structural continuity while suppressing overlap-dominated interface regions. The resulting binary mask was applied to the precomputed (*G, S*) images (computed from the original, unmodified FLIM decays), ensuring that phasor coordinates reflected the original temporal measurements while spatial support was restricted by MSSR-derived separability. Representative MSSR parameters used in the implemented pipeline included sigma_px = 2.43 (PSF-width estimate in pixels), order = 1 (MSSR^1^), width = 0, mesh = 0, ftI = 0, amplification = 0, and intNorm = False. A representative masking threshold applied to the 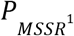 support map in experimental analyses was mean_min = 0.04.

### Sparrow-limit Gaussian emitter simulations for MSSR-derived probability-support mapping

To illustrate how MSSR improves separability at the Sparrow criterion using the bounded intermediate output 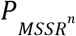, synthetic data was generated from two equal-intensity Gaussian emitters and processed in one and two spatial dimensions. Two emitters were modeled as identical Gaussian intensity distributions with standard deviation σ. In one dimension, the individual contributions were defined as:

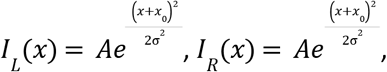

with equal amplitudes *A*. The composite raw signal was:

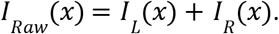

The emitter separation was set to the Sparrow limit for equal-width Gaussians by imposing a center-to-center distance of 2σ, implemented as *x* = σ. The horizontal axis was expressed in units of σ by plotting the dimensionless coordinate 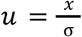 . For visualization, profiles were normalized by their maximum value:

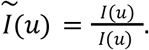

Signals were evaluated on a uniform grid spanning [*u*_*min*_, *u*_*max*_] with *u*_*min*_ =− 4 and *u*_*max*_ = 4, corresponding to [ − 4σ, 4 σ]. A fixed sampling step Δ*u* was used to ensure smooth profiles. The same axis definition was used to generate one-dimensional profiles and to define the coordinate system for the two-dimensional fields.

MSSR was applied as an intensity-domain operator to enhance separability in the synthetic signals without altering any time-resolved quantity. MSSR was computed on the normalized raw intensity field (one-dimensional or two-dimensional). Two MSSR orders were evaluated: MSSR^0^ (first-stage mean-shift transformation output) and MSSR^1^ (one additional mean-shift iteration). For downstream masking, the workflow used the intermediate MSSR output obtained prior to the final intensity remapping step. This intermediate output was treated as a probability-like spatial support map in [0,1] and denoted 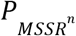, with order-specific forms:

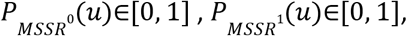

for one-dimensional signals, and analogously 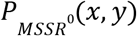 and 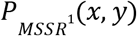 for two-dimensional fields. The bounded normalization was implemented by rescaling the intermediate MSSR output to the unit interval. Binary support masks were constructed by thresholding the probability-like MSSR outputs:

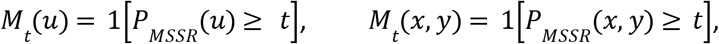

where 1[·] denotes the indicator function and *t* ∈ [0, 1] is the threshold level. Threshold selection was explored using quartile-like cutoffs *t* ∈ [0, 0. 25, 0. 50, 0. 75], where *t* = 0 corresponds to an unthresholded condition *M*_0_ = 1 everywhere.

Two-dimensional emitter fields were constructed as the sum of two isotropic Gaussian spots:

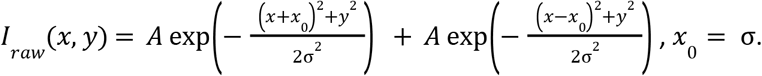

The field was sampled on the normalized coordinate system 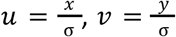 and normalized to unit maximum prior to MSSR processing. MSSR^0^ and MSSR^1^ were computed on this normalized field to obtain 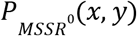 and 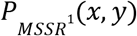. For each threshold *t* ∈ [0, 0. 25, 0. 50, 0. 75], masked probability-density maps were generated as:

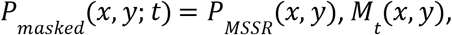

and displayed using a fixed color scale spanning 0–1. Simulations and rendering were performed in Python using numpy, scipy, and matplotlib. MSSR processing used napari_superres, and the intermediate probability-like output *P*_*MSSR*_ was extracted prior to the final intensity remapping step and rescaled to [0,1] for thresholding and visualization.

### Simulation framework for TD-FLIM and FD-FLIM signals and phasor-coordinate validation

Synthetic simulations were used to validate phasor behavior for monoexponential versus multiexponential decays and to illustrate equivalence between TD-derived and FD-derived phasors under matched modulation frequency and calibration assumptions. Simulations were implemented in Python using numpy and scipy and generated one-dimensional temporal signals only.

Global simulation settings were parameterized by modulation frequency f and angular frequency \omega:

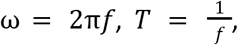

where *T* is the modulation period. A representative FD condition used *f* = 80*MHz*, corresponding to *T* = 12. 5*ns*. Signals were discretized into samples = 256 points over one period, yielding *t* ∈ [0, *T*]. Signal scaling used mean = 10.0 (arbitrary units) and background = 0.0. An instrument-response term was included as a narrow, impulse-like response controlled by a phase-location parameter (zero_phase) and an optional width parameter (zero_stdev); a figure-generating configuration used zero_stdev = None to produce a narrow response.

Time-domain decay synthesis used monoexponentials:

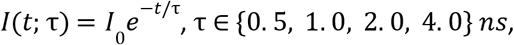

and multiexponential decays as amplitude-weighted mixtures:

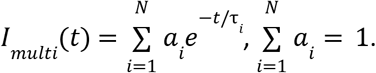

A representative two-component mixture used equal fractions (fractions = [0.5, 0.5]), and a single-component reference decay was generated for calibration and validation.

Frequency-domain analogs were generated over one period and rendered on a phase axis ϕ in degrees derived from time *t* by:

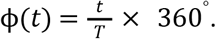

Composite FD signals were generated by linear combination of component contributions consistent with the amplitude fractions used in the TD mixtures.

Phasor coordinates were computed from lifetimes using the monoexponential mapping:

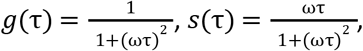

and mixture phasors were computed as amplitude-weighted averages:

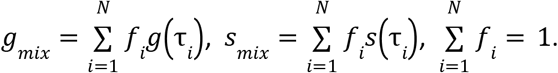

Internal validation compared phasors computed from simulated signals (signal-to-phasor transform) against theoretical lifetime-derived phasors after reference-based calibration. A representative reference lifetime used for calibration was reference_lifetime = 0.3 ns. Agreement was enforced by numerical tolerance checks (numpy.testing.assert_allclose).

### In silico FLIM model for two non-interacting fluorophores with controlled spatial overlap

To quantify temporal mixing under PSF overlap and to test MSSR-based mitigation under fully parameterized conditions, a two-emitter in silico model was implemented as repository-tracked notebooks. The simulation workflow used PhasorPy for FLIM-signal synthesis and phasor transforms, numpy for array construction, scipy (notably scipy.signal) for convolution, and matplotlib for visualization. MSSR operations were executed through napari_superres.

Two independent fluorophores were modeled as monoexponential emitters with lifetimes τ_1_ = 1*ns* and τ_2_= 4*ns*. Temporal sampling was defined by a modulation frequency *f* = 80*MHz*, with ω = 2π*f* and laser period *T* = 1/*f* = 12. 5*ns*. Temporal waveforms were discretized with N=256 samples per period/binning window. The baseline intensity scale used mean = 1e4 and background = 0. A narrow, impulse-like instrument-response proxy was included by specifying zero_phase = 0.08 and zero_stdev = None, and phasor calibration used a reference lifetime reference_lifetime = 0.06 ns.

Time-resolved fluorescence signals were synthesized using PhasorPy’s lifetime-to-signal simulator for single-component decays and a two-component mixture with equal fractional contributions (fractions = [0.5, 0.5]) to validate amplitude-weighted linearity in phasor space. Each fluorophore temporal signal *Ik*(*t*) was assigned to a single pixel location (*i*_*k*_, *j*_*k*_), encoding temporal information at spatially discrete emitter coordinates prior to optical blurring.

A synthetic FLIM image stack *F*(*y, x, t*) was constructed with spatial dimensions x_dim = 127, y_dim = 127 and temporal dimension t_dim = 256. Emitters were placed on the central row (cy = center = 63) with controlled horizontal separations expressed in units of the PSF standard deviation σ. Spatial sampling used pxsz_nm = 5 nm per pixel and an isotropic Gaussian PSF with sigma_nm = 100 nm, giving σ *px* = σ *nm* /*pxsz nm* = 20 pixels and 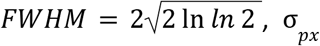, σ *px*. Separation conditions included complete overlap and separations spanning the range from sub-resolution to well-resolved, including a minimum-separation condition at 1.6σ implemented as 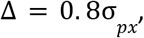, and additional separations such as 2.0σ, 2.5σ, 3.0σ, 4.0σ, and 6.0σ via Δ = {1. 0, 1. 25, 1. 5, 2. 0, 3. 0}σ *px*.

Optical blurring was modeled by convolving each time slice with the Gaussian PSF. For each time index *t*,

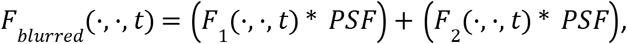

where * denotes 2D convolution computed using FFT-based convolution (scipy.signal.fftconvolve, mode=“same”). This produced a time-resolved image stack in which temporal information remained fluorophore-specific at the source level but became spatially mixed through PSF overlap.

Per-pixel phasor components were computed by Fourier decomposition along the temporal axis using PhasorPy:

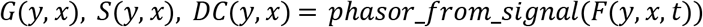

with the temporal axis explicitly set (axis = 2). Systematic offsets were corrected by reference-based calibration (phasor_calibrate) using τ_*ref*_ = 0. 06*ns* at *f* = 80*MHz*. Background suppression was implemented by intensity thresholding on the mean component using phasor_threshold with mean_min = 0.75, yielding thresholded maps *DC*_*th*_, *G*_*th*_, *S*_*th*_ for lifetime extraction and visualization.

Phase and modulation were computed as:

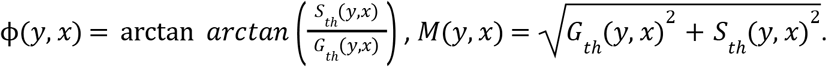

Apparent phase and modulation lifetimes were computed using Weber identities:

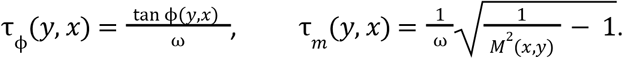

To stabilize calculations in low-amplitude regimes, lower limits were imposed as mod_lower_limit = 0.36 and phase_lower_limit = 0.1 prior to lifetime mapping; multiplicative scaling factors were unity (factor_tm = 1, factor_tp = 1), and optional correction terms were disabled (mod_correction = False, phase_correction = False) to preserve correspondence with analytic phasor expressions.

For quantitative summaries, lifetimes were sampled along the central row and at three landmark pixels: the two emitter-coordinate pixels (*x*_1_, *y*_*c*_) and (*x*_2_, *y*_*c*_), and the midpoint (*x*_*c*_, *y*_*c*_) (saddle/reference pixel). These values were tracked across separations to quantify the dependence of recovered apparent lifetimes on spatial overlap.

Phasor-space segmentation for simulated data used polar cursors defined by phase and modulation bounds, implemented with mask_from_polar_cursor. Three classes were encoded via:

- phase_min = [0.78, 0, 0.453], phase_max = [1.13, 0, 0.555]
- modulation_min = [0.438, 0, 0.77], modulation_max = [0.568, 0, 0.905]

Cursor overlays were rendered using PhasorPlot.polar_cursor, and reciprocity mapping to image space was visualized with pseudo_color, using the thresholded mean intensity as the support image.

### MSSR-based mitigation of temporal mixing in simulated FLIM data

MSSR was applied as an intensity-domain operation to improve spatial separability while maintaining the original temporal signal definitions of each fluorophore. For unresolved or near-unresolved separations, MSSR processing was performed on the mean intensity projection *DC*(*y, x*) obtained from the phasor decomposition (equivalently, the sum over the temporal axis). MSSR was executed via napari_superres using a mean-shift formulation that concentrates local intensity density toward maxima. The simulation implementation used order = 1 (MSSR^1^), fwhm = 2.4 (PSF width parameter in the MSSR interface), and disabled auxiliary options (amp = 0, mesh = 0, ftI = 0, intNorm = 0).

The MSSR-processed intensity map was converted into a photon-redistribution field using intensity2photons, and temporal information was reassigned in a two-fluorophore setting using temporal_reassignment_2F. This reassignment preserves the per-fluorophore temporal waveforms *I*_1_ (*t*) and *I*_2_ (*t*) while altering the spatial allocation of their contributions according to the MSSR-derived redistribution, yielding a reassigned stack *F*_*MS*_ (*y, x, t*). Phasor computation, reference calibration, intensity thresholding (mean_min = 0.75), and lifetime extraction (τ_*M*_, τ_ϕ_) were then repeated on *F*_*MS*_ (*y, x, t*) using the same modulation frequency (80 MHz), the same Weber identities, and the same numerical lower limits (mod_lower_limit = 0.36, phase_lower_limit = 0.1). Comparisons between pre- and post-MSSR simulations were performed at the level of mean intensity structure, phasor distributions in (G,S) space, apparent lifetime maps, and landmark-pixel lifetime values at emitter coordinates and midpoint, enabling attribution of changes in lifetime mixing to spatial reassignment rather than to changes in the decay model.

### Three-component TD-FLIM phasor analysis and MSSR masking

A three-component cellular FLIM dataset was analyzed in phasor space to evaluate MSSR-mediated suppression of Temporal Blurring in regions of multi-structure spatial overlap. The specimen collected with a Leica TCS SP8 STED 3X platform equipped with a FALCON-FLIM module (photon-counting hybrid detectors, galvanometric scanning, and a 63× high-numerical-aperture oil-immersion objective), corresponds to the same fixed-cell multicolor FLIM slide used throughout this work, including the DAPI channel for three-color TD-FLIM acquisitions; each spectral channel was processed independently using the same phasor formalism. The region of interest (ROI) reported in Figure 9 was selected in the diffraction-limited image to include spatial overlap among nucleus (DAPI), microtubules (AF555), and mitochondria (AF532), thereby maximizing the prevalence of mixed pixels for downstream evaluation.

Phasor coordinates were computed directly in the processing notebook using the PhasorPy (phasorpy) Python library. Time-tagged photon data were imported from PicoQuant ‘.ptu’ files and organized as time-resolved image stacks with axes (x, y, t), where t indexes the TCSPC histogram bins over the 80 MHz excitation period. Pixel-wise phasors (G, S) and the mean intensity (DC component) were obtained by Fourier decomposition at the selected analysis harmonic using the PhasorPy implementation, yielding spatial maps of DC, G, and S and a corresponding phasor distribution in (G, S) space.

Instrument-related phase shifts and timing offsets were corrected by reference calibration using a homogeneous lifetime standard acquired under matched acquisition settings. A homogeneous solution of ATTO 488 was used as the calibration reference. The reference phasor was computed from this standard within the same PhasorPy workflow and used to calibrate experimental phasors by aligning the measured reference phasor to the theoretical phasor expected for the ATTO 488 reference lifetime at the 80 MHz modulation frequency. The resulting calibration transform was applied to the three-component cellular dataset prior to phasor visualization, cursor-based segmentation, and MSSR-masked re-analysis.

To suppress background and low-count pixels, intensity thresholding was applied to the DC image to generate a binary support mask that was propagated to the G and S maps prior to visualization and segmentation. Spatial stabilization of the phasor maps was performed by median filtering of G and S using a small kernel (3 × 3) with repeated application, consistent with the denoising procedure described in the Methods. Three-component behavior was assessed by the emergence of a triangular phasor distribution. Endpoint populations were identified by cursor selection near each vertex of the triangle, and reciprocity mapping was used to project phasor selections back into image space for visualization of the spatial distribution of each component and mixed regions.

For MSSR-assisted mitigation of temporal mixing, Mean-Shift Super-Resolution (MSSR^1^) was applied to the mean-intensity image to generate an MSSR-derived probability-like map. The MSSR output was rescaled to [0, 1], and a fixed threshold (8% in Figure 9) was applied to create a binary MSSR mask that restricts analysis to high-likelihood emitter regions prior to phasor transformation and segmentation. The MSSR mask was applied to the ROI dataset, and phasor distributions were recomputed from the masked pixels to evaluate changes in cluster compactness and mixing signatures. Cluster compactness was quantified using an ellipse-area ratio, defined as the area of a covariance-based confidence ellipse fitted to a cluster after MSSR masking divided by the area before masking. Centroid displacement was quantified as the Euclidean distance between pre- and post-masking cluster centroids in (G, S) space. These metrics were used to confirm reduced phasor dispersion (reduced mixing) together with centroid stability (preservation of intrinsic decay signatures).

### Software, libraries, and computational environment

Experimental data processing, visualization, phasor operations, filtering, MSSR masking, and model fitting were performed in Python using numpy, pandas, scipy (including scipy.ndimage and scipy.optimize), matplotlib, tifffile, PhasorPy (phasorpy), and napari_superres. Simulation plotting additionally used cmcrameri for colormap selection. Phasor coordinate computation from time-domain decays was performed in LifetimeXplorer v1.10.9 with GPU acceleration when available, exporting mean intensity and (g,s) matrices for Python-side processing.

## Data and code availability

Analysis scripts and notebooks used to reproduce MSSR-FLIM processing, simulations, and quantitative metrics are available in the project repository: https://github.com/MarioGoG/MSSR-FLIM-2025.git. All images analyzed are available via the Zenodo repository and can be retrieved from the DOI provided here: “Two- and Three-color confocal FLIM images of U2OS Gatta Quant samples” https://doi.org/10.5281/zenodo.19028067

## Acknowledgments

MGG acknowledges the team members of the Laboratorio Nacional de Microscopía Avanzada for their valuable feedback during the preparation of this manuscript. We are also deeply grateful to the members of the PhasorPy Project for developing a powerful and accessible tool that was instrumental in building the *in silico* model used in this study. We are also thankful to the BINA, GBI, and LABI initiatives for providing funding opportunities to attend courses and workshops.

## In memoriam

MGG dedicates this work to the memory of Philippe Bastiaens (1963–2025), whose pioneering work on fluorescence lifetime imaging and the concept of temporal blurring helped inspire the theoretical perspective developed in this study.

---

**Fig. S1.**
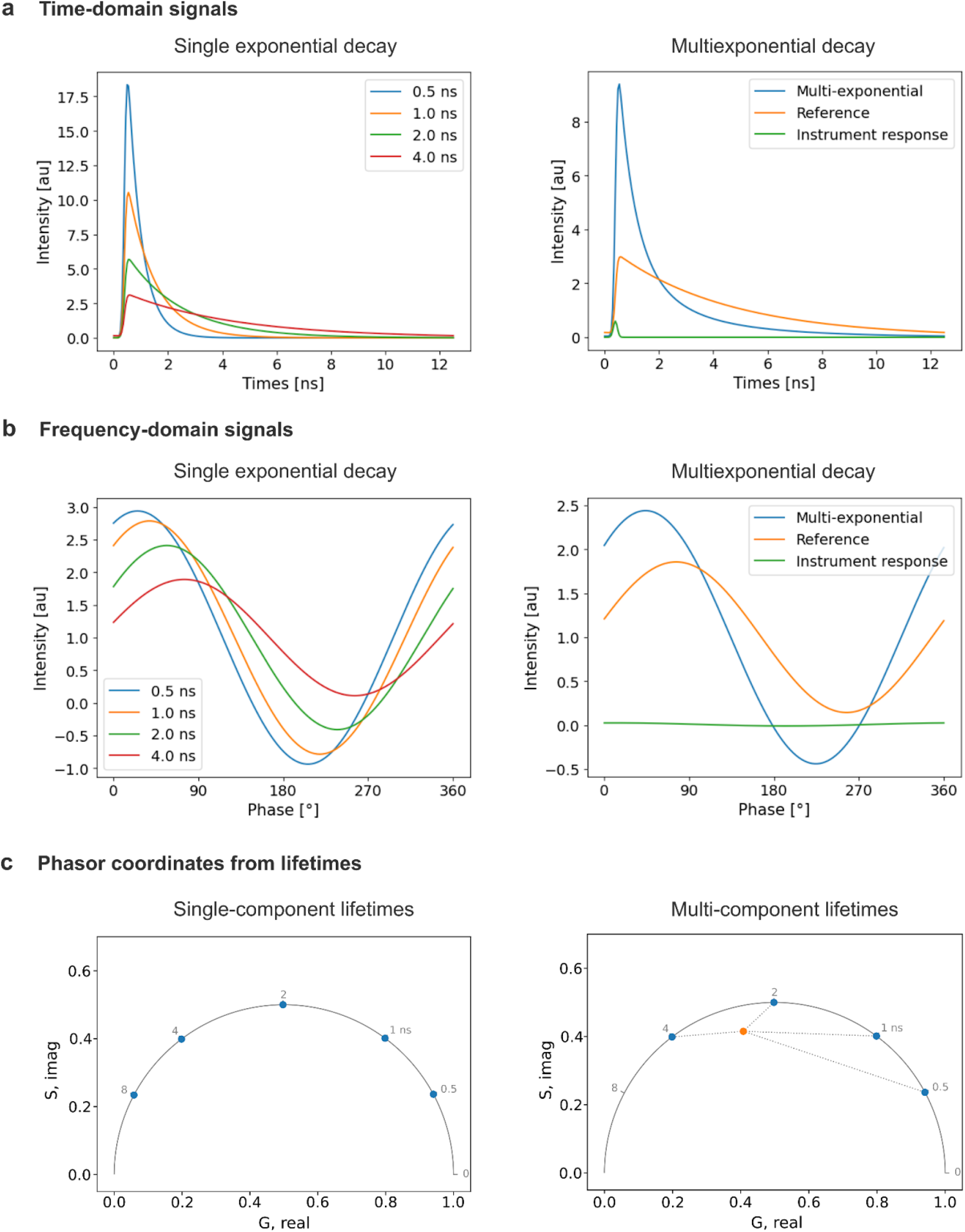
Conceptual link between TD-FLIM, FD-FLIM, and phasor mixing for monoexponential versus multicomponent decays. (a) Time-domain signals. Left: Ideal monoexponential decays with lifetimes of 0.5 ns (blue), 1.0 ns (orange), 2.0 ns (green), and 4.0 ns (red). Right: A representative multicomponent (composite) decay (blue) shown alongside a reference decay used for calibration (orange) and the instrument response function, IRF (green). (b) Frequency-domain signals. Left: Frequency-domain emission waveforms for the same monoexponential lifetimes, using the same color code as in (a). Increasing lifetime produces increased phase delay and reduced modulation. Right: Frequency-domain representation of a mixed (multicomponent) signal (blue), shown alongside the reference waveform (orange) and IRF-related contribution (green). (c) Phasor coordinates derived from the signals in (a) and (b). For a given analysis harmonic (and after appropriate calibration/IRF handling), TD-FLIM and FD-FLIM yield equivalent phasor coordinates (G,S), and therefore provide indistinguishable phasor plots in the ideal case. Left: Monoexponential lifetimes map to single points on the semicircle (labels indicate lifetime ordering). Right: A multicomponent decay maps inside the semicircle (orange point), consistent with linear combination of lifetime components. Colors in panels (a,b) index lifetime components, whereas the orange color in panel (c, right) denotes the multicomponent phasor. Dotted segments are geometric guides illustrating that the mixed phasor lies within the convex region spanned by the contributing monoexponential phasors; fractional contributions are encoded by the interior point position rather than by the segment lengths.

**Fig. S2.**
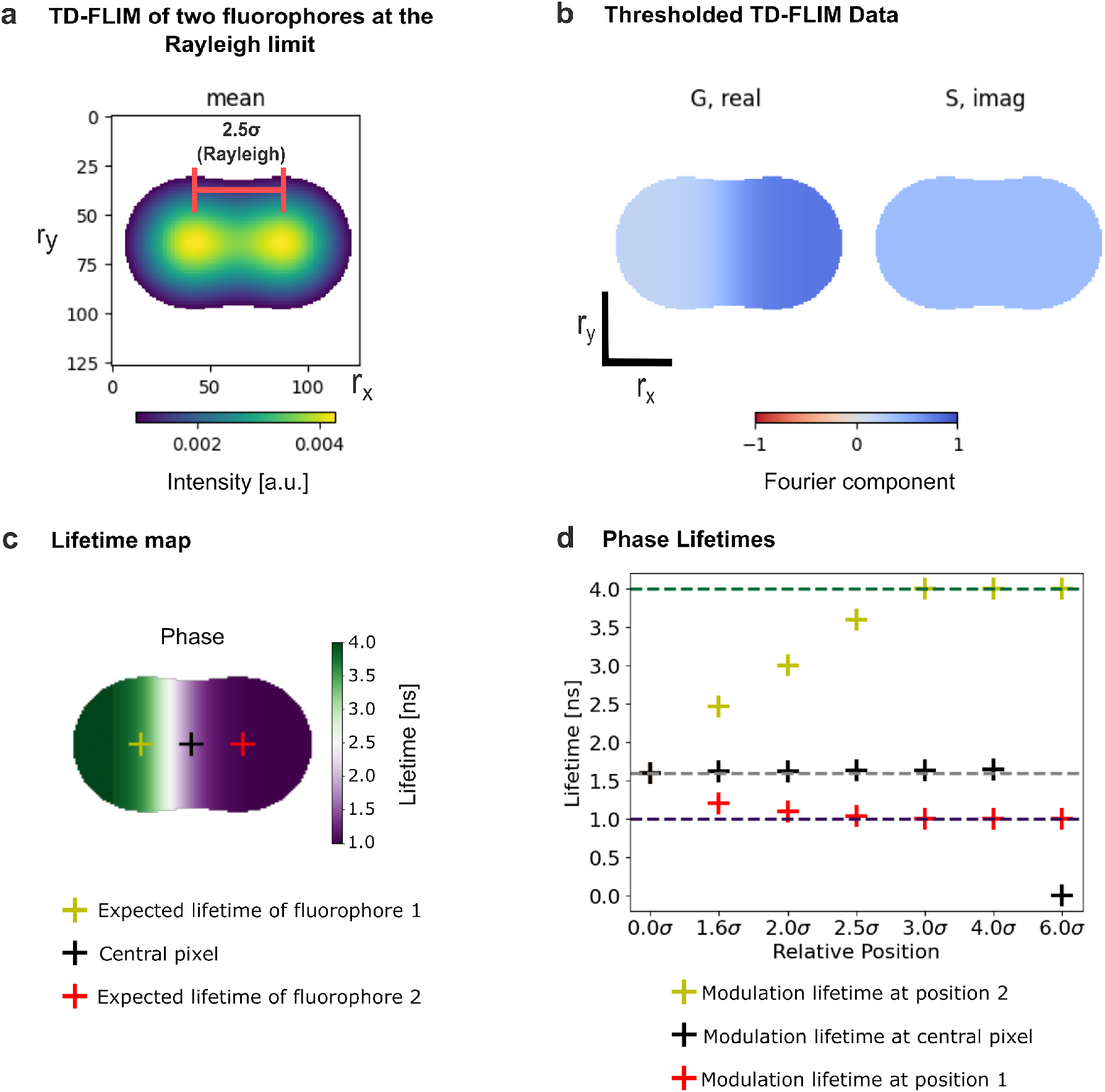
Phase lifetime transition shift in the PSF-overlap regime. Simulated TD-FLIM dataset for two independent, non-interacting fluorophores positioned at 2.5σ. The phase lifetime map is displayed alongside the corresponding spatial intensity reference. Panels (a–c) show simulated TD-FLIM images of two fluorophores positioned at 2.5σ. (a) Steady-state fluorescence intensity (mean from Fourier analysis) with intensity thresholding applied to suppress background. Axes represent pixel coordinates (*r*_*x*_, *r*_*y*_). (b) Phasor component maps: G (real) and S (imaginary). (c) Phase lifetime map computed from Fourier components. The transition zone in phase lifetime is shifted toward the long-lifetime fluorophore relative to the geometric midpoint of the two emitter positions, consistent with preferential capture of phase information under PSF-induced mixing. (d) Phase lifetimes sampled at the centers of each fluorophore position and at the central reference pixel, plotted as a function of inter-fluorophore separation (σ).

**Fig. S3.**
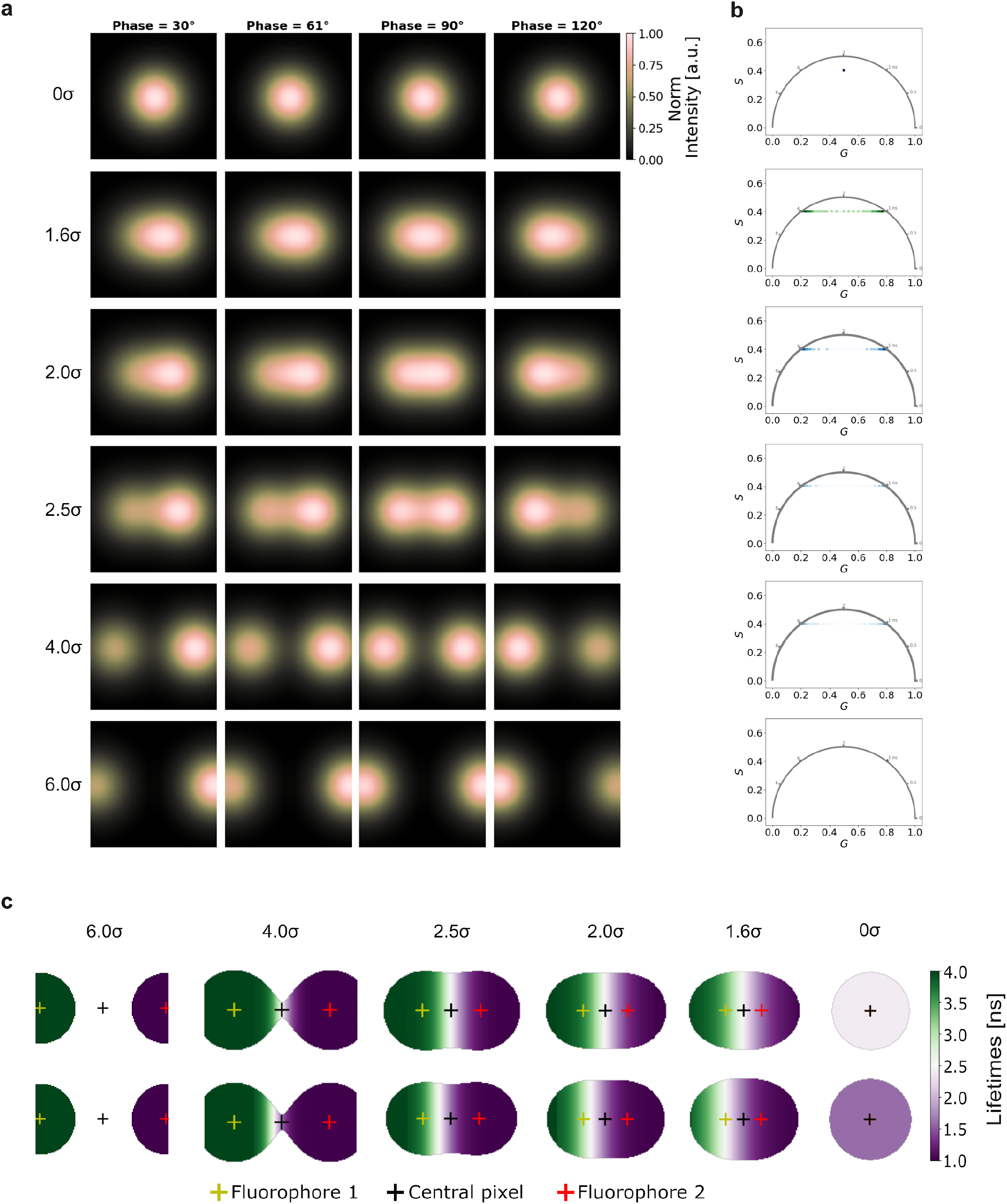
FD-FLIM *in silico* simulation of Temporal Blending induced by PSF overlap. Frequency-domain implementation of a two-fluorophore *in silico* model consisting of two independent, non-interacting and monoexponential emitters with lifetimes of 4ns (left) and 1ns (right).(a) Representative sinusoidally modulated gaussian-blurred intensity frames at four relative phase angles (30°, 60°, 90°, and 120°). Each row corresponds to a different inter-fluorophore separation expressed in units of the Gaussian PSF standard deviation σ, ranging from complete overlap (0σ) to well-separated emitters (6σ). As the separation decreases, PSF-induced spatial mixing becomes increasingly pronounced. (b) Corresponding phasor plots derived from the data in panel (a). With decreasing inter-fluorophore distance, phasor clusters progressively broaden and shift toward an amplitude-weighted mixed position inside the universal semicircle, reflecting apparent multi-exponential behavior arising solely from spatial averaging. This demonstrates that TB emerges in FD-FLIM through PSF-mediated photon mixing, even when the underlying fluorophores have strictly monoexponential decay. (c) Modulation- and Phase-lifetime maps associated with the simulated FD-FLIM data, illustrating how spatial overlap produces intermediate apparent lifetimes in regions between emitters. Together, these simulations confirm that TB manifests analogously in FD-FLIM as a geometric-optical effect rather than a change in intrinsic decay kinetics.

**Fig. S4.**
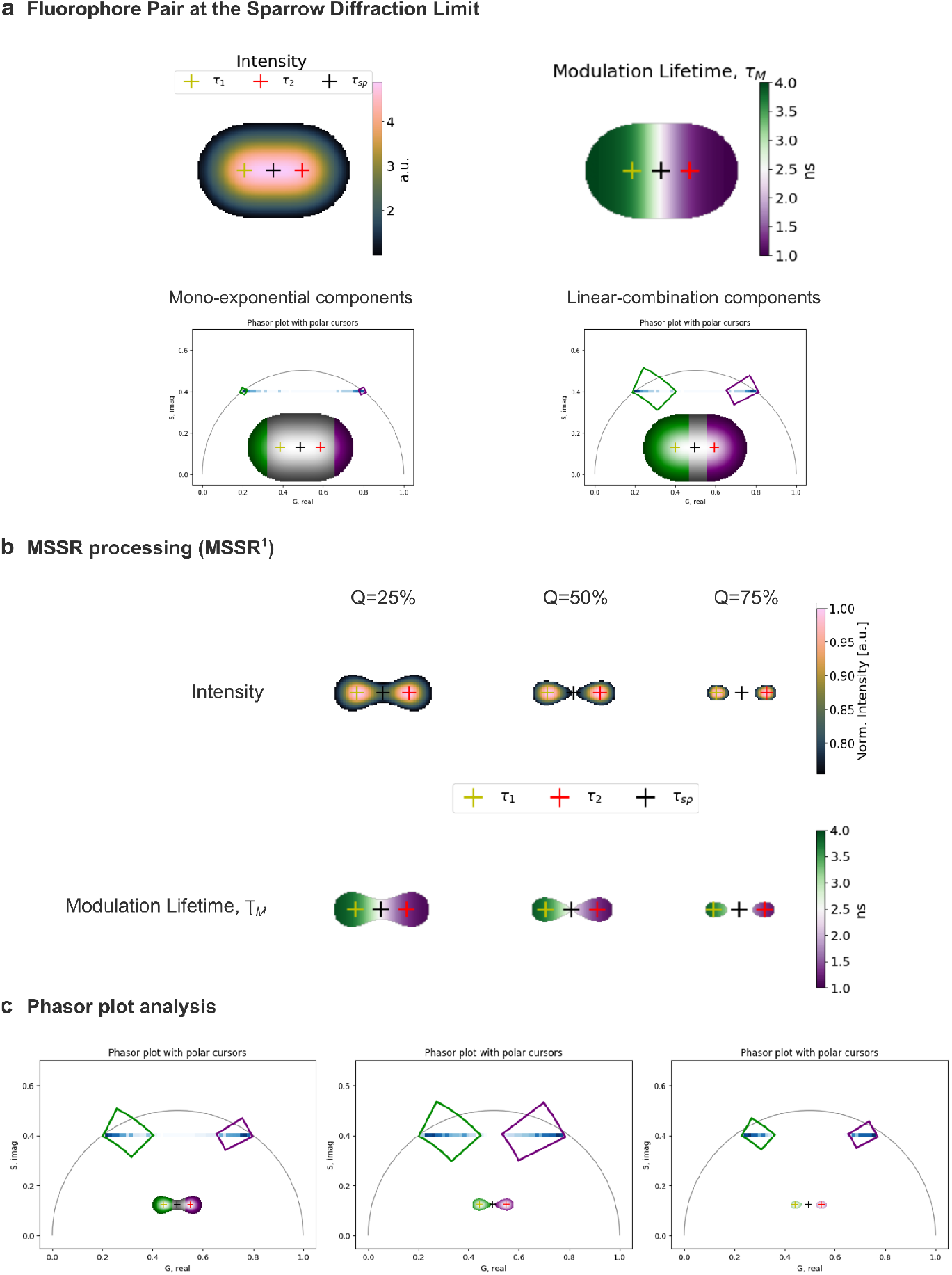
Percentile-based MSSR masking progressively reduces Temporal Blending at the Sparrow limit. Two independent, non-interacting fluorophores simulated at the Sparrow diffraction limit (2.0σ; same dataset as Fig. 8) are analyzed using MSSR-derived spatial probability masks. (a) Diffraction-limited intensity image, corresponding modulation lifetime map, and phasor plots illustrating monoexponential components (lying on the universal semicircle) and linearly combined components arising from PSF-induced overlap, the insets show selected pixel regions on the modulation lifetime maps that were segmented in the phasor space by the rectangle-polar cursors to illustrate with the reciprocity principle the extension difference of the monoexponential pixels and the linearly-combined pixels. (b) MSSR^1^-processed intensity images thresholded at the 25th, 50th, and 75th percentiles (top row), with the corresponding modulation lifetime maps shown below. Increasing the percentile threshold progressively restricts the analysis to spatial regions with a higher probability of originating from individual emitters, thereby reducing effective PSF overlap. (c) Phasor analysis of the MSSR-masked datasets reveals a monotonic reduction in phasor cluster spread and a depletion of intermediate apparent lifetimes as the threshold increases, while the monoexponential phasor positions remain stable. Together, these results demonstrate that percentile-based MSSR masking systematically mitigates Temporal Blending by spatially excluding mixed pixels without altering the intrinsic fluorescence decay signatures.

