## Supporting Information for "PSF-Driven Spatio-Temporal Blending in Fluorescence Lifetime Imaging Microscopy and Its Mitigation via Mean-Shift Super-Resolution-Based Masking"

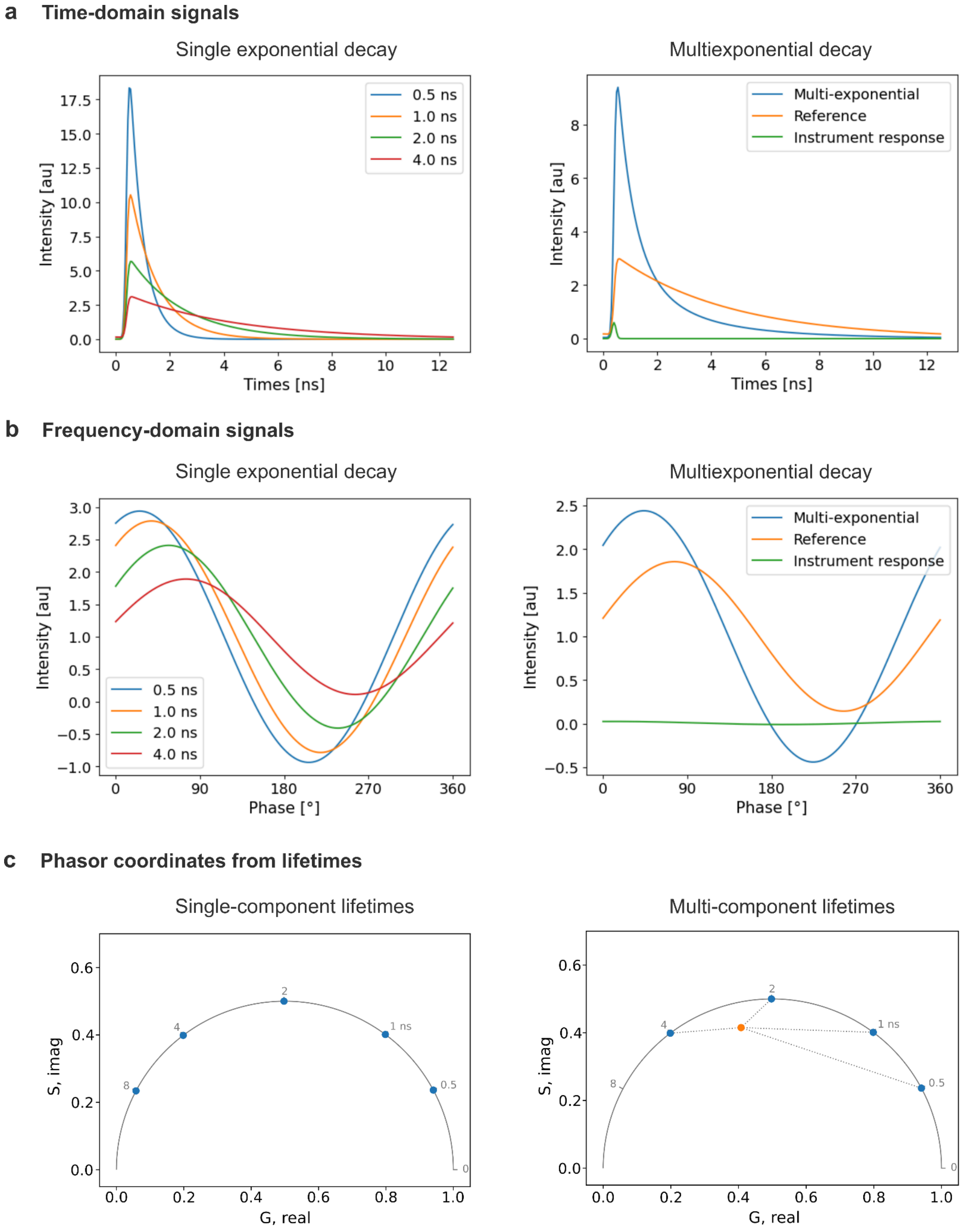


**Fig. S1 Overview of TD-FLIM and FD-FLIM with the phasor analysis of the single-exponential cases vs the multi-exponential cases (Molecular meaning of a linear combination of lifetimes).** a) Plots showing different intensity decay profiles form a Time-Domain perspective in which the intensity is exponentially decreasing. Left: Four monoexponential fluorescence decays ranging from 0.5 ns to 4 ns. The reference signal used for calibration and the Instrument Response Function (IRF) are not shown for simplicity. Right: The blue plot represents a linear combination of all the monoexponential profiles in the left plot yielding an amplitude-weighted single decay profile, the graph was computed taking into account each amplitude weight of 25%. b) Plots showing the time-varying intensity profiles from a

Frequency-Domain perspective. In this case the signal is measured relative to a modulated excitation signal (not shown for simplicity) and the modulation of the signal is plotted against the phase dimension (horizontal axis). Left: Four monoexponential fluorescence oscillations ranging from 0.5 ns to 4 ns. Notice how they gradually shift towards larger phase angles, thus providing information about their phase lifetime, whereas their modulations are smaller with increasing lifetimes. Reference signal and IRF are not shown for simplicity. Right: Plots showing the mixed FD signals into one single sinusoidal signal. c) Phasor plots for the lifetime profiles in a) and b). Both perspectives TD- and FD-FLIM are equivalent due to the Fourier Transform that relates Frequency with Time. Right: The monoexponential profiles appear in the phasor space as single points along the semicircumference positioned anticlockwise from smaller to larger lifetimes. Left: The Orange point inside the semicircumference represents the linear combination of all the lifetime profiles involved. the distances from that point to each lifetime point within the semicircumference measures the fraction of each fluorescence lifetime participating in the resulting signal.

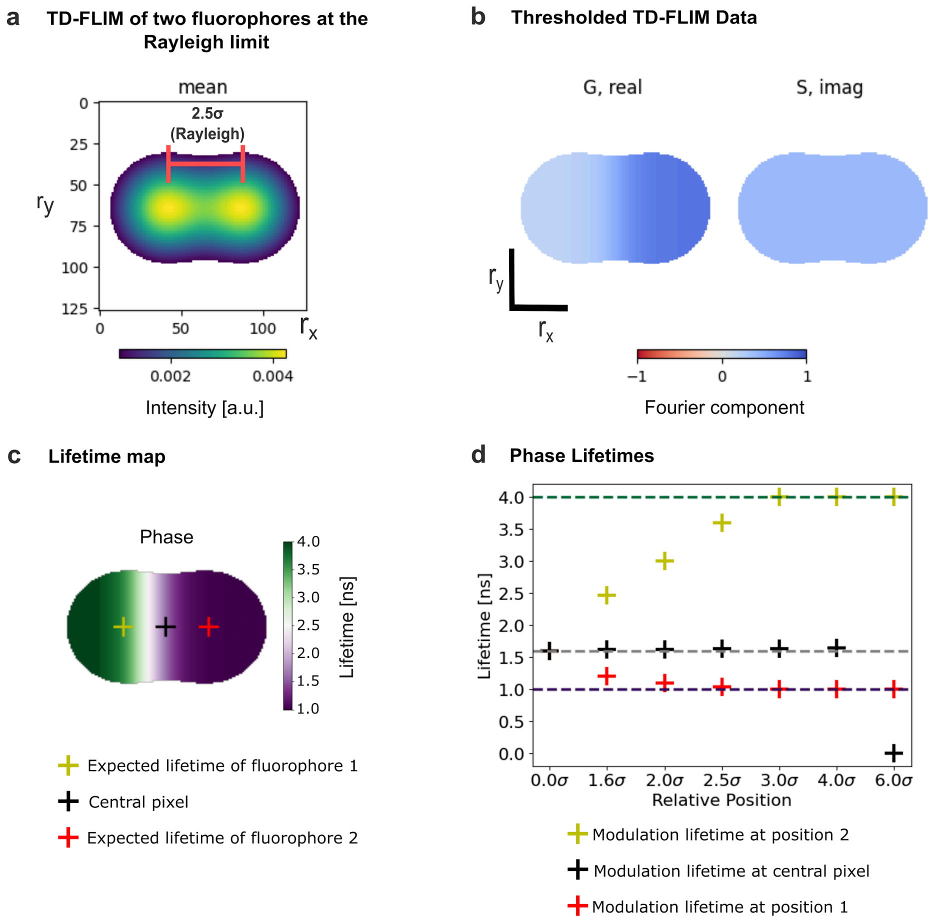


**Fig. S2. Phase lifetime transition shift in the PSF-overlap regime. Simulated TD-FLIM dataset for two independent, non-interacting fluorophores positioned at 2.5σ.** The phase lifetime map is displayed alongside the corresponding spatial intensity reference. Panels (a–c) show simulated TD-FLIM images of two fluorophores positioned at 2.5σ. (a) Steady-state fluorescence intensity (mean from Fourier analysis) with intensity thresholding applied to suppress background. Axes represent pixel coordinates $(r_x, r_y)$. (b) Phasor component maps: G (real) and S (imaginary). (c) Phase lifetime map computed from Fourier components. The transition zone in phase lifetime is shifted toward the long-lifetime fluorophore relative to the geometric midpoint of the two emitter positions, consistent with preferential capture of phase information under PSF-induced mixing. (d) Phase lifetimes sampled at the centers of each fluorophore position and at the central reference pixel, plotted as a function of inter-fluorophore separation (σ).

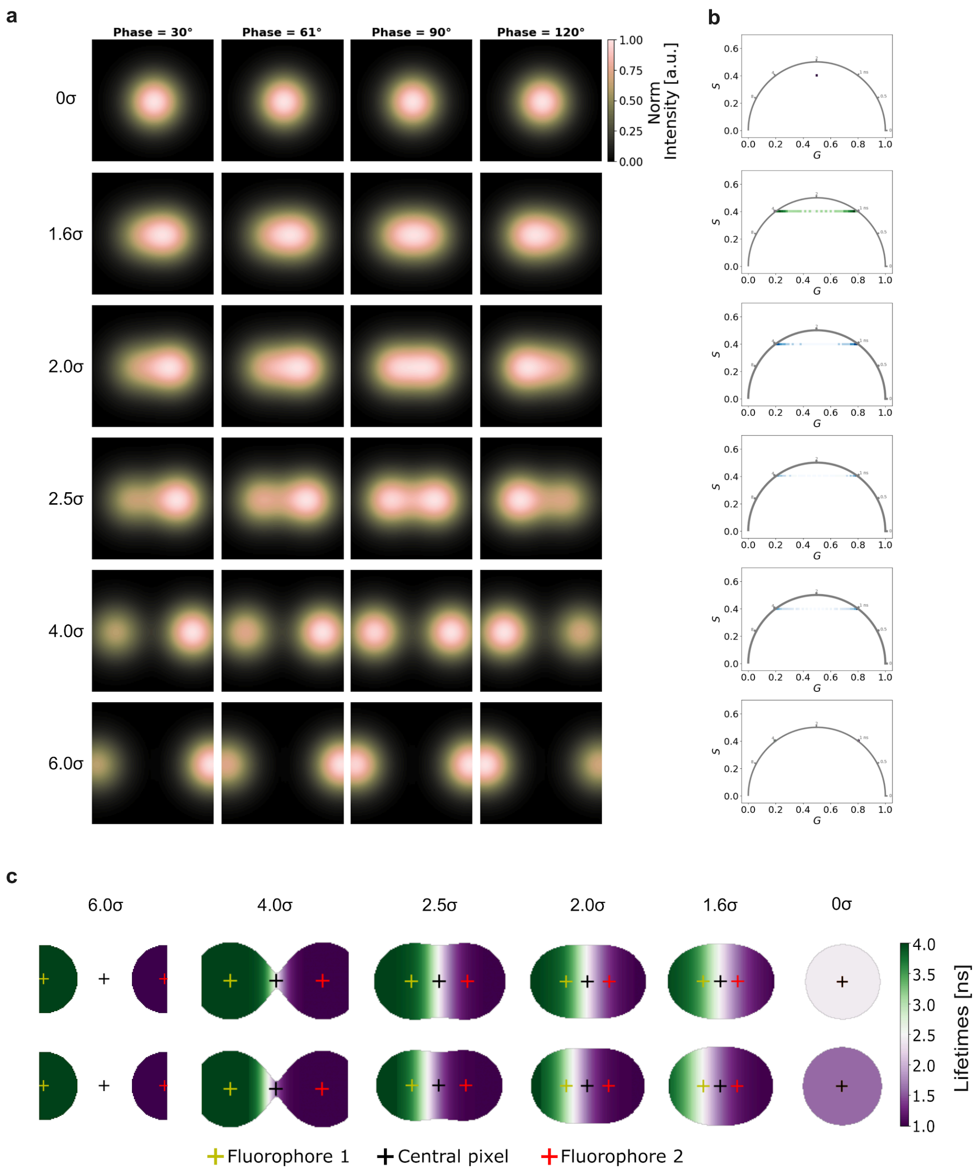


**Fig. S3. FD-FLIM *in silico* simulation of Temporal Blending induced by PSF overlap.** Frequency-domain implementation of a two-fluorophore *in silico* model consisting of two independent, non-interacting and monoexponential emitters with lifetimes of 4ns (left) and 1ns (right).(a) Representative sinusoidally modulated gaussian-blurred intensity frames at four relative phase angles (30°, 60°, 90°, and 120°). Each row corresponds to a different inter-fluorophore separation expressed in units of the Gaussian PSF standard deviation σ, ranging from complete overlap (0σ) to well-separated emitters (6σ). As the separation decreases, PSF-induced spatial mixing becomes increasingly pronounced. (b) Corresponding phasor plots derived from the data in panel (a). With decreasing inter-fluorophore distance, phasor clusters progressively broaden and shift toward an amplitude-weighted mixed position inside the universal semicircle, reflecting apparent multi-exponential behavior arising solely from spatial averaging. This demonstrates that TB emerges in FD-FLIM through PSF-mediated photon mixing, even when the underlying fluorophores have strictly monoexponential decay. (c) Modulation- and Phase-lifetime maps associated with the simulated FD-FLIM data, illustrating how spatial overlap produces intermediate apparent lifetimes in regions between emitters. Together, these simulations confirm that TB manifests analogously in FD-FLIM as a geometric-optical effect rather than a change in intrinsic decay kinetics.

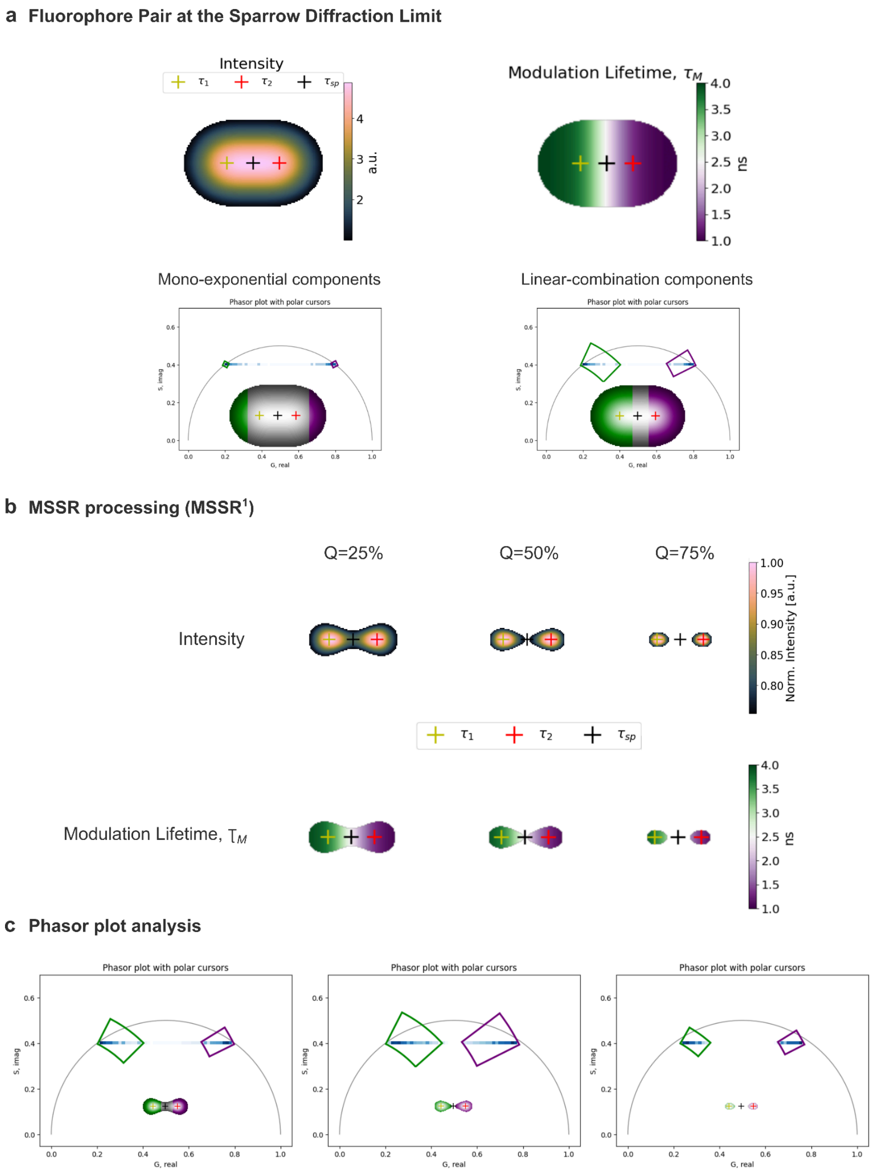


**Fig. S4. Percentile-based MSSR masking progressively reduces Temporal Blending at the Sparrow limit**. Two independent, non-interacting fluorophores simulated at the Sparrow diffraction limit (2.0σ; same dataset as Fig. 8) are analyzed using MSSR-derived spatial probability masks. (a) Diffraction-limited intensity image, corresponding modulation lifetime map, and phasor plots illustrating monoexponential components (lying on the universal semicircle) and linearly combined components arising from PSF-induced overlap, the insets show selected pixel regions on the modulation lifetime maps that were segmented in the phasor space by the rectangle-polar cursors to illustrate with the reciprocity principle the extension difference of the monoexponential pixels and the linearly-combined pixels. (b) MSSR[1]-processed intensity images thresholded at the 25th, 50th, and 75th percentiles (top row), with the corresponding modulation lifetime maps shown below. Increasing the percentile threshold progressively restricts the analysis to spatial regions with a higher probability of originating from individual emitters, thereby reducing effective PSF overlap. (c) Phasor analysis of the MSSR-masked datasets reveals a monotonic reduction in phasor cluster spread and a depletion of intermediate apparent lifetimes as the threshold increases, while the monoexponential phasor positions remain stable. Together, these results demonstrate that percentile-based MSSR masking systematically mitigates Temporal Blending by spatially excluding mixed pixels without altering the intrinsic fluorescence decay signatures.

#### Phasor Cloud Metrics

To quantify the effect of MSSR on the phasor cloud, we computed several statistical descriptors from the 2D distributions of the real ($G$) and imaginary ($S$) components. Pixels below the applied intensity threshold were excluded (NaN values ignored).

| **Metric** | **Equation** |
|---|---|
| Centroid | $(\bar{G}, \bar{S}) = (\frac{1}{N}\sum_{i=1}^{N} G_i, \frac{1}{N}\sum_{i=1}^{N} S_i)$ |
| Centroid displacement | $\Delta r = \sqrt{(\bar{G}_{after} - \bar{G}_{before})^2 + (\bar{S}_{after} - \bar{S}_{before})^2}$ |
| Ellipse area (95% confidence) | $A = ab\pi$, <br> $a = \sqrt{\lambda_1 \chi_\alpha^2}, b = \sqrt{\lambda_2 \chi_\alpha^2}; \lambda_1, \lambda_2 = eig(\Sigma); \lambda_1 \geq \lambda_2$ <br> $\chi_\alpha^2 = \chi^2_{0.95, 2} = 5.991$ |
| Ellipse area ratio | $R_A = \frac{A_{after}}{A_{before}}$ |

Interpretation table of metrics

| **Metric** | **Expected Values / Behavior** | **Interpretation** |
|---|---|---|
| Centroid displacement ($\Delta r$) | ≈ 0 (no displacement) | Preservation of intrinsic lifetimes. Large displacements (>0.05 in normalized phasor space) indicate distortion of decay kinetics information. |
| Ellipse area ratio ($A_{after}/A_{before}$) | < 1 | Ratio < 1: reduced phasor cloud spread. <br><br> Ratio ≈ 1: no effect. <br><br> Ratio > 1: increased uncertainty or artifact introduction. |

These metrics evaluate the performance of MSSR when enhancing the precision of lifetime estimation in phasor space without introducing artifacts. The centroid stability demonstrates preservation of intrinsic lifetime values, while the reduction in ellipse area and increase in anisotropy indicate suppressed temporal blurring and improved definition of the mixing axis. The covariance and orientation analyses further support that MSSR selectively reduces noise without biasing the underlying decay relationships. Together, these results provide quantitative evidence that MSSR improves both spatial and temporal fidelity in FLIM phasor analysis.

### Phasor-Based Representation of FLIM Signals Under Spatial Diffraction (IRF Neglected)

1. Scope and conceptual motivation

Fluorescence Lifetime Imaging Microscopy (FLIM) measures the temporal decay of fluorescence intensity at each pixel of an image. In practice, the signal recorded by a detector pixel does not originate from a single point in space but from a finite three-dimensional volume determined by the optical point spread function (PSF) of the microscope.

The phasor representation of FLIM maps fluorescence decays into a two-dimensional complex plane using Fourier components of the temporal signal. This representation provides a compact and model-free way to visualize lifetime populations and mixtures.

Here, we outline a conceptual and mathematical framework that explicitly incorporates spatial diffraction into the phasor formulation. The goal is not to introduce a complete theory but rather to provide a mechanistic interpretation clarifying how spatial convolution imposed by the PSF can influence the phasor distributions observed in FLIM images.

For clarity, the derivation neglects the temporal instrumental response function (IRF). Under this assumption, deviations from monoexponential phasor behavior arise from spatial signal mixing rather than temporal broadening introduced by the detection system.

2. Phasor representation of fluorescence decays

For a fluorescence decay measured at a spatial position $r$ , the intensity as a function of time is denoted $I(r, t)$.

The normalized phasor components at modulation frequency $\omega$ are

$$g(r, \omega) = \frac{\int I(r,t) cos(\omega t) dt}{\int I(r,t) dt}, \; s(r, \omega) = \frac{\int I(r,t) sin\,(\omega t)\, dt}{\int I(r,t) dt}$$

These define the complex phasor

$$G(r, \omega) \; = \; g(r, \omega) \; + \; i\, s(r, \omega)$$

Using Euler's identity the same quantity can be written

$$G(r, \omega) \; = \; \frac{\int I(r,t) e^{i\omega t}\, dt}{\int I(r,t)\, dt}$$

which highlights that the phasor corresponds to the normalized first harmonic of the Fourier transform of the fluorescence decay.

3. Monoexponential decays and the universal semicircle

For a monoexponential decay $I(t) = Ae^{-t/\tau}$, the theoretical phasor becomes

$$G(\omega) = \frac{1}{1+i\omega\tau}$$

Separating real and imaginary parts yields

$$g(\omega) = \; \frac{1}{1 + (\omega\tau)^2}, \; s(\omega) = \; \frac{\omega\tau}{1 + (\omega\tau)^2}$$

All monoexponential decays therefore lie on the universal semicircle of the phasor plot, centered at (0.5,0) with radius 0.5. This geometric property forms the basis of the phasor approach for identifying lifetime populations and mixtures.

4. Molecular mixtures within a voxel
If several fluorescent species coexist within the same spatial location, the decay can be written

$$I(r,t) = \Sigma A_k(r)\, e^{-t/\tau_k}$$

Defining fractional intensity contributions $f_k(r)$ , the resulting phasor becomes

$$G_{mix}(r,\omega) = \Sigma f_k(r) G_k(\omega)$$

Thus, the phasor of a molecular mixture is a convex combination of the phasors of the individual species. In the case of two components the measured phasor lies along the straight line connecting the two monoexponential phasors. This behavior reflects intrinsic biochemical heterogeneity and is independent of optical resolution.

5. Spatial diffraction and optical signal formation
In a real microscope, each pixel collects photons from a finite spatial region defined by the PSF. The measured signal at detector position r0 can therefore be written as a spatial convolution

$$I_{meas}\left(r_0, t\right) = \int PSF\left(r_0 - r\right) I_{true}(r,t) dr$$

where $I_{true}(r,t)$ represents the local fluorescence decay. Physically, this expression reflects that photons detected at a given pixel originate from a weighted neighborhood determined by the optical response of the system.

6. Phasor of the measured signal
The measured phasor at pixel $r_0$ is defined as

$$G_{meas}\left(r_0, \omega\right) = \frac{\int I_{meas}\left(r_0, t\right) e^{i\omega t}\, dt}{\int I_{meas}\left(r_0, t\right) dt}$$

Substituting the convolution expression yields a double integral involving both space and time.

7. Exchange of spatial and temporal integrals
Because the PSF is independent of time and fluorescence intensities are non-negative, the order of integration can be exchanged according to the Fubini–Tonelli theorem. This allows separation of spatial and temporal operations and leads to an expression where the Fourier transform of the decay is computed locally before spatial averaging.

First, let us define the local integrated intensity as

$$I_0(r) = \int I_{true}(r,t)\, dt$$

and the local phasor

$$G_{true}(r, \omega) = \frac{\int I_{true}(r,t)e^{i\omega t}\, dt}{I_0(r)}$$

The measured phasor $G_{meas}\left(r_0, \omega\right)$ is defined by substituting the $I_{meas}\left(r_0, t\right)$ value from subsection 5.

$$G_{meas}\left(r_0, \omega\right) = \frac{\int [\int PSF\left(r_0 - r\right) I_{true}(r,t) dr] e^{i\omega t} dt}{\int [\int PSF\left(r_0 - r\right) I_{true}(r,t) dr] e^{i\omega t} dt}$$

As explained at the beginning of this subsection, we rearrange the order of the integration according to the Fubini-Tonelli theorem, and because the PSF is independent of time, this yields:

$$G_{meas}\left(r_0, \omega\right) = \frac{\int [PSF\left(r_0 - r\right) \int I_{true}(r,t) e^{i\omega t} dt] dr}{\int [PSF\left(r_0 - r\right) \int I_{true}(r,t) dt] dr},$$

Finally, substituting the expressions for the local integrated intensity and the local phasor results in:

$$G_{meas}\left(r_0, \omega\right) = \frac{\int PSF\left(r_0 - r\right) I_0(r) G_{true}(r,\omega) dr}{\int PSF\left(r_0 - r\right) I_0(r) dr}$$

8. Interpretation as a weighted spatial average

The measured phasor can therefore be expressed as

$$G_{meas}\left(r_0, \omega\right) = \int w(r) G_{true}(r, \omega) dr$$

with weights

$$w(r) = \frac{PSF\left(r_0 - r\right) I_0(r)}{\int PSF\left(r_0 - r\right) I_0(r) dr}$$

Where $w(r)$ satisfies

$$\int w(r) dr = 1$$

Meaning these weights are non-negative and normalized to unity. Consequently, the measured phasor corresponds to a convex spatial average of the underlying local phasors.

9. Conceptual contrast with FMCE

This spatial-averaging interpretation differs conceptually from the Frequency Modulation Capture Effect (FMCE) model proposed for frequency-domain FLIM. FMCE attributes lifetime dominance to amplitude-driven frequency capture in the temporal domain. In contrast, the framework outlined here emphasizes a geometric-optical origin of apparent lifetime mixing through spatial convolution of overlapping PSFs before phasor transformation.

10. Takeaway statement

Under the assumptions adopted here (neglecting IRF and considering diffraction-limited detection), the phasor measured at a detector pixel can be interpreted as a PSF-weighted spatial average of local phasors. This observation provides a possible mechanistic explanation for intermediate phasor populations appearing at structural interfaces in FLIM images, even when individual emitters exhibit monoexponential decay.

11. Cautionary note

The formulation presented here should be regarded as a conceptual model aimed at clarifying the role of spatial diffraction in phasor-space representations of FLIM data. Further theoretical analysis and experimental validation will be required to determine the generality of this interpretation across imaging modalities and biological systems.

12. Experimental Signatures of Spatial Mixing

1. Resolution Dependence
   Varying pinhole size or numerical aperture changes PSF width and thus alters $G_{meas}$. In contrast, intrinsic molecular mixtures remain invariant.
2. ROI Scaling
   Increasing the ROI area shifts $G_{meas}$toward the mean phasor of included regions—characteristic of spatial averaging.
3. Blurred Lifetime Transitions
   Lifetime maps exhibit intermediate lifetimes at edges of heterogeneity due to overlapping PSFs.
4. Straight-Line Trajectories
   In phasor plots, pixels at interfaces lie along straight lines connecting neighboring pure lifetimes.

13. Consequences for Lifetime Interpretation

Spatial diffraction leads to:

- Apparent multiexponential decays where none exist,
- Underestimation of lifetime contrast between adjacent domains,
- Loss of spatial precision in heterogeneous samples,
- Intermediate phasor positions inside the semicircle.

These artifacts do not reflect real biochemical interactions or energy transfer, but rather optical averaging imposed by the PSF.

14. Mitigation Strategies

- Improve optical confinement: smaller pinhole (confocal), higher NA objectives, multiphoton excitation, or super-resolution approaches (e.g., STED-FLIM, MSSR-FLIM).
- Spatial segmentation: restrict analysis to homogeneous subregions.
- Phasor unmixing: define pure phasors (endmembers) and project mixed points along connecting lines to estimate spatial fractions.
- Resolution tests: compare images acquired with different PSF volumes; spatial mixing will manifest as resolution-dependent phasor shifts.
